# Multi-omic screening identifies RBMXL3 as a primate-specific RNA-binding protein and candidate regulator of RNA metabolism in human spermatogenesis

**DOI:** 10.1101/2025.09.23.677621

**Authors:** Agata J. Barszcz, Katarzyna Tutak, Joanna Zyprych-Walczak, Erik Dassi, Erkut Ilaslan, Małgorzata Dąbrowska, Agnieszka Malcher, Marta Olszewska, Dominik Cysewski, Agnieszka Nadel, Tomasz Kolanowski, Zofia E. Madeja, Michał Hrab, Alexander Yatsenko, Maciej Kurpisz, Zbigniew Warkocki, Natalia Rozwadowska

## Abstract

**Background:** RBMXL3 is a primate-specific gene localized on the X chromosome, which expression is detectable mainly in the male gonad. So far, very little is known about the RBMXL3 protein function and its molecular interactions. However, recent reports mention the *RBMXL3* gene in the context of human spermatogenesis, cancer, and a breathing disorder that affects newborns. In this study, we investigate the RBMXL3’s molecular network on a genome-wide scale using the human seminoma cell line (TCam-2) as a male germline *in vitro* model.

**Methods:** By using transcriptomic (RNA sequencing (RNA-seq) and enhanced crosslinking and immunoprecipitation (eCLIP)) and proteomic (Co-immunoprecipitation coupled with Mass Spectrometry, (Co-IP-MS)) approaches we show RBMXL3 importance in RNA metabolism. Additionally, Western Blot, qRT-PCR, immunostaining, and confocal imaging were used in order to investigate the function of RBMXL3. Finally, we used a plasmid-based L1 retrotransposition assay to demonstrate the suppressive effect of RBMXL3 on human Long Interspersed Nuclear Element-1 (LINE-1, L1) retrotransposition.

**Results:** Our RNA-seq data show that RBMXL3 expression drives gene expression changes and influence alternative splicing in human cells. Moreover, by performing eCLIP we provide a proof that RBMXL3 binds to a wide range of RNA transcripts. Additionally, we confirmed the nuclear localization of RBMXL3 in TCam-2 cells and its presence in spermatogonia and spermatocytes within the human testis. Finally, we report for the first time that RBMXL3 restricts human LINE-1 retrotransposition.

**Conclusions:** Our findings for the first time identify primate-specific RBMXL3 protein as a new upstream regulator of RNA metabolism, characterized by broad RNA-binding activity in human TCam-2 cells. Finally, we show that RBMXL3 expression heavily reduces LINE-1 retrotransposition in human cells, underlying RBMXL3 importance in maintaining genome integrity. Our data suggest that RBMXL3 may contribute to the regulation of transcriptome dynamics in male germ cells, while its broader functional implications remain to be determined.

**Graphical abstract:** 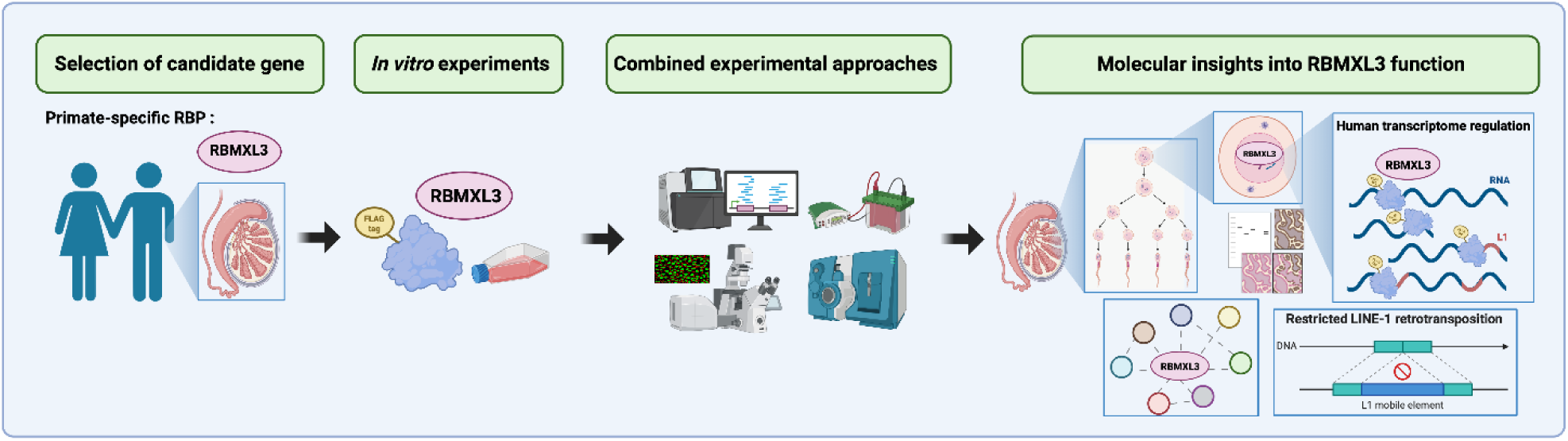

Created with BioRender.com

## Background

Infertility has become a global health issue that affects millions of couples worldwide [1–4]. Since approximately half of infertility cases are associated with male factor it is necessary to explore the mechanisms underlying spermatozoa differentiation. A plethora of factors must cooperate to ensure successful male gametes differentiation during spermatogenesis, and the complexity of this process makes it error-prone. Unfortunately, our knowledge about the genetic regulation of human germ cell development and spermatogenesis is fragmented. High-throughput sequencing technologies, including whole-exome sequencing (WES) and whole-genome sequencing (WGS) studies, pave the way for seeking new candidate genes that may be responsible for some fertility disorders. Recent genome-wide genetic screening of non-obstructive azoospermia (NOA) patients identified novel candidate genes, possibly associated with spermatogenesis failure [5]. Among the potential NOA-associated gene candidates, the *RBMXL3 (RNA-binding motif X-linked-like-3)* gene, coding for a primate-specific protein, was identified. *RBMXL3* is located on Xq23 within the first intron sequence of *LRCH2*. The *RBMXL3* is a primate-specific, intronless gene which is transcribed in the opposite direction to its host gene and encodes the RBMXL3 protein consisting of 1067 amino acids (114.9 kDa). This protein shares 50 % identity with its ancestral protein RBMX (RNA binding motif protein X-linked, also localized on X chromosome). *RBMXL3* originated, as many other *RBMX*-like genes, from spliced *RBMX* mRNA transcript in the process called retrotransposition [6–9]. The RBMX protein, previously called heterogeneous nuclear ribonucleoprotein G – hnRNP G, is ubiquitously expressed in multiple tissues and contains an N-terminal RRM (RNA recognition motif) [6]. In addition to the RRM domain, the RBMXL3 protein contains a unique long C-terminal glycine-rich region with not yet defined function [10]. A schematic comparison of RBMXL3 and related RBMX proteins has been previously published by Liu et al [10].

Multiple lines of evidence support the involvement of the RBMX-family proteins in post-transcriptional regulation of gene expression [11], including mRNA splicing [12]. RBMXL2 controls splicing patterns in meiotic prophase during spermatogenesis, and *Rbmxl2* gene knock-out results in male mice sterility [13]. Also RBMX is involved in splicing and associates co-transcriptionally with RNA polymerase II (RNAPII) in humans [14–16]. Importantly, DNA sequence variants in the *RBMX* gene have already been associated with NOA in humans [17].

Alternative splicing regulation has been shown to be especially important in the context of developmental processes and is particularly prevalent in the testis and the brain [18–20]. There is an evident relation between the human brain dysfunctions and testis disorders [21, 22]. For example, the already mentioned RBMX has been linked to Shashi X-linked intellectual disability syndrome characterized, among others, with macroorchidism (abnormally large testes) [23, 24]. Complementarily, studies on a zebrafish *rbmx* orthologue have shown that it is essential for normal brain development [25]. Furthermore, the RBMX protein has been shown to contribute to sister chromatid cohesion whereby its knockdown causes abnormal nuclear morphology [26, 27]. RBMX also contributes to the efficient DNA repair and ensures genome stability during replication [28–30] which is essential during gametogenesis.

According to the Human Protein Atlas (HPA), *RBMXL3* is expressed mostly in spermatogonia and spermatocytes in the human testis. At a lower level, however, its expression has been also found in brain [31]. Mass spectrometry-based proteome-wide studies of healthy human tissues confirmed RBMXL3 protein in the testis and lung [32]. However, wide-scale experiments on human tissues have shown discrepancies concerning the expression of *RBMXL3*, at the mRNA and protein level, indicating RBMXL3 elusive nature [32–34]. Rare variants of *RBMXL3* have been associated not only with NOA [5] but also with *MYCN* non-amplified neuroblastoma, which is the most common extracranial solid tumor in infants and children [35]. Two other reports mention this gene in the context of metastatic colorectal cancer [36, 37]. The only report that showed the first functional aspect of RBMXL3 refers to its presence in the human lung cancer cell line (A549) wherein it was proposed to stabilize the transcript encoding pulmonary surfactant protein B [10].

Since modeling human spermatogenesis *in vitro* is limited, we decided to address RBMXL3 functionality in male germ cells using the TCam-2 cell line, derived from human seminoma (germ cell tumor) [38–41]. In terms of gene expression, these cells resemble human primordial germ cells [42, 43] which are embryonic precursors of male germ cell lineage and hence are a valuable tool for studying germline-specific processes. Here, we generated modified TCam-2 cell lines expressing either RBMXL3-FLAG or GFP-FLAG and used them to characterize transcriptomic and proteomic changes resulting from *RBMXL3* overexpression. By using RNA-seq and eCLIP, we show that RBMXL3 is a promiscuous RNA-binding protein influencing both gene expression as well as splicing of multiple target RNAs. In addition, we identify RBMXL3 protein interactors of which most are involved in post-transcriptional regulation of gene expression, including splicing. Finally, we observe RBMXL3 binding to LINE-1 (L1, Long Interspersed Nuclear Element-1) retrotransposon RNAs and show reduced retrotransposition of L1 in an *in vitro* L1 retrotransposition assay. This study adds important new information on the functionality of primate-specific RBMXL3 placing it as a possible regulator of human spermatogenesis and differentiation processes.

## Methods

### Cell line

The human testicular cancer cell line (TCam-2) was gifted from Prof. Sohei Kitazawa (Division of Molecular Pathology, Kobe University, Japan) [38, 39]. TCam-2 cells were cultured at 37°C in a humidified 5% CO_2_ incubator in Roswell Park Memorial Institute 1640 Medium (RPMI 1640 Medium, GlutaMAX™ Supplement Thermo Fisher Scientific, 61870044) supplemented with 10% heat-inactivated Fetal Bovine Serum (Cytiva, HYCLSV30160.03) and 1% Antibiotic-Antimycotic solution (100X) (Thermo Fisher Scientific, 15240062).

HEK293T cells (used in retrotransposition assay, originally acquired in Invitrogen) were derived from a female. The 293T cells (ATCC Cat# CRL-3216, RRID:CVCL_0063) were obtained from the lab of Jose L. Garcia-Perez (IGMM Edinburgh) [44] and authenticated by SNP genotyping by Eurofins Genomics (Ebersberg Germany), the chromatograms are available online in the supplementary materials to Janecki et al., 2024 [45]. The cells were cultured in monolayers in high glucose Dulbecco’s modified Eagle’s medium (DMEM, Gibco; 41966052) supplemented with 9% (v/v) fetal bovine serum (Gibco; 10270106), 100 U/ml penicillin and 100 µg/ml streptomycin (Merck; P4458) at 37°C in a humidified 5% CO_2_ incubator. Cells were passaged every 2-3 day.

### Generation of TCam-2 cell lines modified by lentiviral transduction

Lentiviral particles were packed using 2^nd^ (for RBMXL3 construct) and 3^rd^ (for eGFP construct) generation packaging system in HEK293T cells. In case of 2^nd^ generation system 1 × 10^6^ HEK293T cells were transfected according to manufacturer’s recommendations (Sigma-Aldrich, CAPHOS) based on calcium phosphate method with the mixture of plasmids: psPAX (Addgene, #12260), MD2G (Addgene, #12259) and pLV[Exp]-Puro-EF1A>hRBMXL3[NM_001145346.1]/3xFLAG (Vector Builder, Vector ID VB180321-1238ccw) in ratio 1:1:2. In case of 3^rd^ generation system following plasmids were used: pRSV-Rev (Addgene, #12253), pMDLg/pRRE (Addgene, #12251), pMSCV-VSV-G (Addgene, #14888), pLV[Exp]-Puro-EF1A>EGFP/3xFLAG (Vector Builder, Vector ID VB180409-1201grs) in ratio 1:1:1:2. After 48 and 72h post transfection pseudoviral particles were collected and filtered through 0.45 μm (Millipore, SLHVR13SL). Viral aliquots were frozen and stored at -80 °C. Next 1 × 10^5^ TCam-2 cells were incubated with the viral supernatant in the presence of polybrene (final concentration 4µg/ml, Sigma-Aldrich, TR-1003-G) for 48h. After incubation with the pseudoviral particles, selection with puromycin (final concentration 1 ug/ml, Thermo Fisher Scientific, A1113803) was initiated and continued for 4 days.

### Transfection of TCam-2 cell line

TCam-2 cells were seeded the day before transfection using JetPrime transfection reagent (Polyplus, 101000046). For immunofluorescence staining experiments, cells were cultured on 15 mm in diameter round cover glasses in a 12-well culture plates, and transfection mixture was prepared using 0.8 μg of plasmid, 1.6 μl of reagent, and 75 μl of buffer, according to the manufacturer’s protocol. Next, cells were transfected using 75 μl of transfection mixture per single well. For other experiments, cells were cultured on 150 mm cell culture dishes and transfection mixture was prepared using 31.45 μg of plasmid, 62.9 μl of reagent, and 1.31 ml of buffer, according to the manufacturer’s protocol. Next, cells were transfected using 1.31 ml of transfection mixture per single dish. The complete medium was changed before the transfection procedure. Cells were cultured in a complete medium with transfection mixtures for 2 days prior to downstream experiments. Transfection efficiency was monitored in our study by fluorescence microscopy 48 h after transfection. using transfection control samples (TCam-2 cells expressing GFP-FLAG). Additionally, all samples were validated via WB and qPCR prior to downstream experiments.

Plasmids used: pLV[Exp]-Puro-EF1A>hRBMXL3[NM_001145346.1]/3xFLAG (Vector Builder, Vector ID VB180321-1238ccw), pLV[Exp]-Puro-EF1A>EGFP/3xFLAG (Vector Builder, Vector ID VB180409-1201grs).

### qRT-PCR and statistical analysis

RNA was isolated with RNeasy Plus Mini Kit (Qiagen, 74136) or AllPrep Mini Kit (Qiagen, 80004) and residual DNA was removed by TURBO DNA-free™ kit (Thermo Fisher Scientific, AM1907) treatment according to the manufacturer’s protocol. One µg of total RNA was reverse transcribed by SuperScript IV Reverse Transcriptase kit (Thermo Fisher Scientific, 18090050) with the use of random primers (Promega, C1181). For quantitative polymerase chain reaction (qPCR) 10 ng of cDNA was mixed with PowerUp SYBR Green Master Mix (Thermo Fisher Scientific, A25778), 0.3 µM of each primer (all primer sequences are listed in Supplemental Table S16) and assayed using CFX96 Touch qPCR System (Bio-Rad). All samples were normalized to ACTB and GAPDH according to the geNorm algorithm. Statistical analyses were done using GraphPad Prism (v8.0.1) and performed using ordinary one-way ANOVA non-parametric test with Tukey’s multiple comparisons test. Data with statistical significance are as indicated, *p< 0.05, **p< 0.01, ***p< 0.001.

### RNA sequencing and bioinformatic analysis

The RNA-seq experiment was done in duplicate and performed using TCam-2 cells transduced with *RBMXL3-FLAG*, *GFP-FLAG* and non-transduced WT. Cells transduced with *GFP-FLAG* and non-transduced WT TCam-2 cells were used as negative controls. The lentiviral transduction was used to generate stable cell lines in order to ensure relatively sustained and uniform expression of proteins across the cell populations, thereby reducing biological variability and increasing the robustness of transcriptomic analyses. Total RNA was isolated from cells and treated with DNase as it was described above. Libraries were prepared using the TruSeq Stranded mRNA LT kit (Illumina), enabling sequencing of polyadenylated mRNAs. Samples were run on the Illumina NovaSeq6000 platform, 150 bp paired-end reads with at least 30 M reads per sample. Library preparation and sequencing were performed by Macrogen NGS Service (Korea).

RNA-seq data pre-processing was performed by removing low-quality reads (Q<30) and sequencing adapters using Trim Galore (v0.6.10) (https://github.com/FelixKrueger/TrimGalore). Further information regarding each specific RNA-seq analysis is provided in the sections below.

### Differential expression analysis of RNA-seq data

The pre-processed RNA-seq reads were mapped to the human reference genome (GRCh38/hg38, Gencode v44) using STAR (v2.7.11a) [46] (--outSAMtype BAM SortedByCoordinate --quantMode GeneCounts). Sorted BAM files were subsequently indexed using samtools (v1.19.2) [47]. Read counts were then normalized by DESeq2 R package [48]. Lowly expressed genes (with less than 0.1 mean CPM in each condition i.e. RBMXL3, GFP or WT) were excluded from further analysis. Next, we performed the differential expression analysis comparing the ‘RBMXL3 vs. GFP’ and the ‘RBMXL3 vs. WT’ conditions using DESeq2. The set of significant differentially expressed genes (DEGs) was determined based on adjusted p-value (FDR) < 0.05 and |log2FC| ≥ 1. As the final list of genes we intersected the significant DEGs of the two comparisons (‘RBMXL3 vs. GFP’, ‘RBMXL3 vs. WT’), and we retrieved only those genes with the same trend (upregulation or downregulation) in both comparisons.

The Gene ontology and pathway enrichment analyses were performed with the topGO and clusterProfiler [49] R packages, using a 0.05 threshold on Benjamini-Hochberg (BH) adjusted p-value.

### Differential splicing analysis of RNA-seq data

To quantify transcripts expression (TPM) in each condition (RBMXL3, GFP and WT) we employed the Salmon tool (v1.10.2) [50] and aligned the pre-processed RNA-seq reads to the reference transcriptome (Gencode v44). The alternative splicing analysis was then performed using the SUPPA2 tool (v2.3) [51, 52]. To identify splicing events a differential splicing analysis was performed setting a threshold of 0.05 on the adjusted p-value and a cutoff of 0.15 on the absolute value of the dPSI. In order to identify potentially more specific splicing events, the final list of differentially spliced genes (DSGs) was prepared by doing an intersection between the significant predicted splicing events from the two comparisons (‘RBMXL3 vs. GFP’ and ‘RBMXL3 vs. WT’). During the SUPPA analysis, the WT and GFP samples were set always as Cond1 and RBMXL3 as Cond2. The Gene ontology and pathway enrichment analyses were performed with the topGO and clusterProfiler [49] R packages, using a 0.05 threshold on BH adjusted p-value.

### L1s differential expression of RNA-seq data

L1s’ expression was quantified using the BAM files obtained from the alignment with STAR and the TEtranscripts tool (v2.2.3) [53]. Counts were generated using the --mode multi option, and the repetitive elements were annotated using the file GRCh38_GENCODE_rmsk_TE.gtf ( https://labshare.cshl.edu/shares/mhammelllab/www-data/TEtranscripts/TE_GTF/). Differential expression analysis of repetitive elements was performed using the DESeq2 R package [48] for two comparisons (‘RBMXL3 vs. WT’ and ‘RBMXL3 vs. GFP’). Specifically, only repetitive elements that belong to the L1 family were considered. Significant L1s were retrieved using a threshold of |*log*_2_*FC*| ≥ 0.585 and the adjusted p-value (FDR) ≤0.05.

### RBMXL3 eCLIP

To obtain high-level protein expression, two biological replicates of each sample, TCam-2 cells transfected with *RBMXL3-FLAG* and *GFP-FLAG* (as control), were used in eCLIP-seq performed by Eclipse BioInnovations (USA). Briefly, 48h after transfection of TCam-2 cells on multiple 150 mm cell culture dishes, the cells were crosslinked at 254-nm UV with an energy setting of 400 mJoules/cm^2^ (Analytik Jena, CL-3000). Overall, the cells were lysed, RNase I treated, and protein-RNA complexes were immunoprecipitated with anti-FLAG antibody coupled to magnetic beads. Sequencing was performed as SE72 on the NextSeq platform (Illumina). The eCLIP experiment was performed by Eclipse BioInnovations’ standard RBP-eCLIP pipeline as previously described [54]. Raw sequencing reads were trimmed of UMIs and adapters, filtered out of reads that mapped to repeat elements, aligned to the reference genome (hg38) with STAR, and filtered of PCR duplicates.

Clusters (regions within the IP sample that contain a pileup of reads) were identified using CLIPper (https://github.com/YeoLab/clipper) [55, 56]. Each cluster was normalized against the paired input sample. Final peaks for each sample were filtered for enrichment over the input (|log2FC| ≥ 2) and statistical significance (adjusted p-value ≤ 0.01). Replicated peaks were then identified by positionally intersecting peaks of each replicate of the same condition with the intersect command of bedtools[57], with the -s option. Aspecific peaks (i.e. those also found in GFP samples) were then filtered out from RBMXL3_FLAG peaks by positional intersection in the same way. Peaks were annotated using the GenomeRanges [58] and wavClusteR R packages [59], while Gene ontology (GO) and pathway enrichment analyses were performed with the topGO and clusterProfiler packages [49], using a 0.05 threshold on BH adjusted p-value.

De novo motif analysis was conducted by using the HOMER findMotifsGenome command [60] and the - len 4,5,6,8,10,12 -size given -rna parameters.

Reads mapping to repetitive elements were then used to compute the enrichment of repetitive elements in eCLIP reads with respect to input samples, obtaining fold-enrichment and entropy values as a measure of confidence.

### Co-immunoprecipitation coupled with mass spectrometry (Co-IP-MS)

3 × 10^6^ *RBMXL3-FLAG* transfected or wild-type (WT, as negative control) TCam-2 cells were lysed with 1ml of RIPA buffer (Sigma-Aldrich, R0278) in the presence of protease inhibitors (Thermo Fisher Scientific, 87786) at 4°C for 30 min, on rotator. Next, lysate was clarified by centrifugation: 4°C, 10 min, 11,000 x g. One ml of total protein lysates was incubated with 40µl of anti-FLAG M2 magnetic beads (Sigma-Aldrich, M8823) at 4°C for 3 h with gentle rotation. After IP reaction, beads were washed three times with ice-cold TBS washing buffer (50 mM Tris HCl, 150 mM NaCl, pH 7.4) and one time with DPBS (Thermo Fisher Scientific, 14190169). IP reactions were performed in three technical replicates. If samples were sent to mass spectrometry (MS), beads were snap frozen with liquid nitrogen and subsequently proceeded to MS analysis. MS analysis was performed in the Mass Spectrometry Laboratory, IBB PAS. Dried beads were suspended in 50μl of 100mM ABC. Proteins were reduced and alkylated with TCEP/MMTS, digested overnight with trypsin (Promega, V5111) as described previously[61]. Peptide digest was purified with HLB 96-well plates (Waters, WAT058951), vacuum dry then dissolved in 98% water (J.T.Baker 4218-03)/ 2% MeCN (J.T. Baker HPLC JT9012)/ 0.1%TFA (Supelco, 80457) and analyzed by an online LC-MS/MS (liquid chromatography coupled to tandem mass spectrometry) using a Nano-Acquity (Waters) UPLC system and QExative Orbitrap mass spectrometer (Thermo Fisher Scientific), by applying peptides to precolumn (nanoACQUITY UPLC Trapping Column Waters) using water containing 0.1% formic acid as a mobile phase and then transferred to a nano-column (nanoACQUITY UPLC BEH C18 Column (75 μm inner diameter; 250 mm long; Waters, 186007482)) using an acetonitrile gradient (5–35% AcN in 180 min) in the presence of 0.1% formic acid with the flow rate of 250 nl/min. Three washing runs ensuring lack of cross-contamination from previous samples preceded each analysis. The column outlet was directly coupled to the ion source of the spectrometer working in the regime of data-dependent acquisition MS to MS/MS switch. Peptides were eluted directly to the ion source of the mass spectrometer. Data were acquired in the m/z range of 300–2000. Data were searched using MaxQuant [62] version 1.5.3.0 against a reference human protein sequence database (79 052 entries) downloaded from UniProt. Platform search parameters: a match between runs (match time window, 0,7 min; alignment time, 20 min); enzyme, trypsin/p specific; max missed, 2; minimal peptide length, 7 aa; variable modification, methionine, and proline oxidation; fixed, cysteine alkylation (Methylthio); main search peptide tolerance, 4.5 ppm; protein FDR 1%. Finally, sixteen proteins identified in each RBMXL3 replicate but not present in negative controls were classified as potential RBMXL3 protein partners. Gene ontology analysis among identified proteins was performed in Cytoscape version 3.9.1 using ClueGO plugin version 2.5.9, 4% enrichment cutoff was used meaning that the number of genes should represent at least 4% of total genes in a given process. STRING analysis of protein-protein interaction (PPI) networks was performed using StringApp version 1.7.1, a 0.4 cutoff score was used which means interactions below this value were filtered out.

### Immunofluorescence of TCam-2 cells

After two washes in DPBS (Thermo Fisher Scientific, 14190169) TCam-2 cells were fixed with 4% paraformaldehyde (Boster Bio, AR1068) at 4 °C for 15 min. Next, cells were washed 3 times with DPBS, permeabilized with 0.55% Triton X-100 in DPBS for 30 mins at RT and blocked for 1h at RT in blocking solution (10% of suitable serum derived from the host animal species of the secondary antibody, diluted in 0.55% Triton-X100 in DPBS). Cells were incubated overnight at 4 °C with primary antibodies diluted in the blocking solution. After washing 3 times with DPBS, the cells were incubated with secondary antibodies diluted in 0.55% Triton X-100 in DPBS for 1 h at RT in the dark. After washing 3 times with DPBS, cells were mounted in Fluoroshield with DAPI histology mounting medium (Sigma-Aldrich, F6057).

The following primary antibodies were used: mouse anti-FLAG 1:500 (Sigma-Aldrich, F1804), rabbit anti-DYKDDDDK 1:2000 (GenScript, A00170), rabbit anti-RBMXL3 in serum 1:500 (gift from Prof J. Alcorn [10]), mouse anti-EIF4A3 1:700 (Santa Cruz, sc-365549), rabbit anti-DDX39B 1:50 (Sigma-Aldrich, HPA058450), rabbit anti-DDX39B 1:400 (Proteintech, 14798-1-AP).

The following secondary antibodies were used: goat anti-mouse Alexa Fluor 594 1:500 (Thermo Fisher Scientific, A-11005), goat anti-rabbit Alexa Fluor 488 1:1000 (Abcam, ab150077), donkey anti-mouse Alexa Fluor 488 1:1000 (Thermo Fisher Scientific, A-21202), goat anti-rabbit Alexa Fluor 594 1:500 (Thermo Fisher Scientific, A-11012).

All images were obtained with Leica DMi8 confocal microscope with Leica STELLARIS STED system equipped with Leica HyD photodetectors (Leica Microsystems). We utilized HC PL APO CS2 objective with magnification of 100 and numerical aperture of 1.4 (Leica Microsystems). Data were collected and processed with Leica Application Suite X (LAS X) software (v4.4.0.24861).

### Western blot

TCam-2 cells were lysed on a rotator for 30 min in RIPA buffer (Sigma-Aldrich, R0278) supplemented with protease inhibitors (Thermo Fisher Scientific, 87786), followed by three rounds of sonication at 4°C [20 s on, 30 s off] using Bioruptor Pico (Diagenode, Belgium). Next, protein lysates were centrifuged at 15,000 x g for 10 min at 4°C and supernatants were mixed with 4X Laemmli Sample Buffer (Bio-Rad, 1610747) with 340 mM DTT (Thermo Fisher, R0861) and denatured for 10 min at 95°C. Samples were then separated on the 8% SDS polyacrylamide gels and transferred to nitrocellulose membranes (Bio-Rad, 1620097). Membranes were blocked with 5% non-fat milk in TBS buffer with 0.1% Tween-20 (TBST) and proceeded to overnight incubation at 4 °C with primary antibodies diluted in the blocking solution. Washing steps (3×10min. with TBST) were followed by horseradish peroxidase (HRP)-conjugated secondary antibodies incubation for 1h at RT. Blots were washed twice with T for 10 min and two times in TBS for 10 min. The chemiluminescent signal was detected using the Clarity Western ECL Substrate (Bio-Rad, 1705061) and developed by the ChemiDoc MP Imaging System (Bio-Rad).

Primary antibodies used: mouse anti-FLAG 1:1000 (Sigma Aldrich, F1804), mouse anti-ACTB 1:15 000 (Santa Cruz Biotechnology, sc-81178), mouse anti-GAPDH 1:10 000 (Santa Cruz, sc-365062), rabbit anti-RBMXL3 in serum 1:2000 (a kind gift from Prof J. Alcorn[10]), rabbit anti-RBMXL2 1:400 (Sigma-Aldrich, HPA051842).

Secondary antibodies used: HRP-conjugated anti-mouse 1:10 000 (Santa Cruz Biotechnology, sc-516102), HRP-conjugated anti-rabbit 1:5000 (Abcam, ab6721).

Protein fractionation was performed according to the previously described method [63]. Primary antibodies used for Western Blot after protein fractionation experiment were as follows: mouse anti-TUBB 1:5000 (Abcam, ab131205), mouse anti-SNRNP70 1:5000 (Santa Cruz, sc-390899), rabbit anti-H3C1 1:5000 (Abcam, ab18521).

### Immunohistochemistry of testis tissue samples

For the immunohistochemistry staining, commercial human adult testis paraffin sections were used (Zyagen, HP-401). Slides were incubated in 60°C for one hour. Immunohistochemistry (IHC) experiment was performed using specific RBMXL3 antibody in serum (a kind gift from Prof J. Alcorn [10]). To determine the appropriate antibody dilution, eliminate false positive results, and reduce the background staining, a series of positive control reactions were performed using slides with normal testis cells present in section derived from patient with seminoma. The immunohistochemistry procedure was performed using EnVisionFlex+ visualization system kit (Dako, Agilent Technologies, K8002). Slides were then dewaxed and epitopes were unmasked in PT-Link device (Dako, Agilent Technologies) with the use of EnVisionFlex Target Retrieval Solution Low pH (Dako, Agilent Technologies, K8005). After that, the slides were incubated with primary antibody (dilution 1:2000; incubation 20 min at RT). The antigen-antibody complex was detected using EnVision FLEX+ Rabbit LINKER (Dako, Agilent Technologies, K8009) for 20 min at RT and localized using 3-3′diaminobenzidine (DAB) as chromogen. In the end, the tissue sections were counterstained with hematoxylin, dehydrated in ethanol, incubated in a series of xylenes, and cover-slipped using mounting medium (Dako, Agilent Technologies). The protein expression was evaluated with the use of light microscope (Leica DMi8).

### LINE-1 retrotransposition assay

The assays were performed as described [64]. Briefly, 293T cells were seeded at 150,000 per well into 12-well plates (Greiner Bio-one) in 1ml DMEM high-glucose medium (Invitrogen) without antibiotics. Next day the cells were transfected with 750ng 99_PUR_RPS_EGFP and 200ng of either RBMXL3 encoding plasmid (pLV[Exp]-Puro-EF1A>hRBMXL3[NM_001145346.1]/3xFLAG (Vector Builder, Vector ID VB180321-1238ccw)) or pKK_TEV_MBP (pZW165) [65, 66] using 2,5 μl Lipofectamine2000 (Thermo Fisher Scientific) in a total of 100 μl OPTI-MEM (Thermo Fisher Scientific). Next day the medium was exchanged for one containing penicillin/streptomycin (Sigma) and 1µg/ml puromycin (Invivogen) to select cells which received the 99_PUR_RPS_EGFP plasmid. After 2 days medium was exchanged. After a total time of 4 days of selection the cells were detached by treatment with trypsin and analyzed by flow cytometry using Attune NxT flow cytometer (A24861; Thermo Fisher Scientific). The cells in which retrotransposition had occurred (0,01-2,2% of the entire population depending on the test condition) expressed EGFP and were identified by excitation with a blue laser (λ=488nm) and detection using a bandpass filter (BL1 λ=530/30nm). Other cells did not express EGFP. The live cell population was set by gating FSC and SSC parameters (gate 1), followed by subgating single cells (‘singlets’) using the FSC-A (area) versus FSC-H (peak height) parameters. Control non-transfected (NT) cells were analyzed to set reference background EGFP fluorescence (gating <0,005% of the NT cells). A control transfection with EGFP (pKK_TEV_EGFP, pZW171) and mCherry (pKK_TEV_mCherry, pZW174) [65] encoding plasmids (125ng each per well) and the RBMXL3-FLAG and MBP encoding plasmids were done to test whether RBMXL3-FLAG expression influences EGFP and/or mCherry expression. mCherry was analyzed by excitation with a yellow laser (λ =561nm) and detection with the band pass filter (YL2 λ =620/15nm). No cross-talk with EGFP is observed.

Pools of cells were sampled and used to analyze expression of the RBMXL3-FLAG and MBP proteins. The cell lysates were prepared as described [66] and separated onto Novex 4-20% gradient Tris-glycine gels (Thermo Fisher Scientific), wet-transferred on Protran nitrocellulose membrane (Amersham, 0,45µm) in a buffer supplemented with methanol to 20% [66]. Membranes were stained with ponceau S solution, cut into 3 pieces (upper for staining RBMXL3-FLAG, middle for Lamin B, and lower for MBP or EGFP), blocked in 5% low-fat milk in TBST20 and probed with primary antibodies against either FLAG (M2, Merck, F3165), Lamin B (ProteinTech, 12987-1-AP), MBP (NEB, 8032S), or EGFP (control transfection to assess the impact of RBMXL3-FLAG expression on EGFP production; antibodies sc-9996 from Santa Cruz) in 1:1000 (FLAG), and 1:4000 (other Abs) dilutions in 5% skimmed milk in TBST20 for 16hr at 6°C. Secondary HRP-coupled antibodies (from goat) were against rabbit Abs (111-035-144) or mouse Abs (115-035-146) (both Jackson Immunoscience (0,8mg/ml)), used in 1:20,000 dilutions in milk for 2hr at RT. Signals were obtained by chemiluminescence using Clarity Western ECL Substrate (Bio-Rad) and photographed using Uvitec Q9 Alliance CCD camera.

## Results

### RBMXL3 is a nuclear protein enriched in the chromatin fraction

Due to limited information about RBMXL3 function and lack of available models with native *RBMXL3* expression, we generated a cellular model with transient overexpression of this protein using human seminoma cell line. We transfected TCam-2 wild-type (WT) cells with vector encoding either *RBMXL3* tagged with FLAG or *GFP* tagged with FLAG (used as a control). The expression of *RBMXL3-FLAG* was confirmed on both RNA and protein levels (Fig. 1A,B). To confirm that the signal observed with the anti-FLAG antibody is a fusion with RBMXL3 we used antibodies specific to: (i) RBMXL3 (kindly gifted by Prof. J. Alcorn [10]); and (ii) RBMXL2, which according to the Human Protein Atlas (HPA) detects not only RBMXL2 but also the RBMXL3 protein (Fig. 1C; Supplemental Fig. S1). RBMXL3-FLAG band appeared at ∼140 kDa and larger than expected (∼115 kDa plus ∼3 kDa 3xFLAG_tag). This is in line with a previous study and may result from posttranslational modifications [10]. To investigate RBMXL3 subcellular localization, the cells overexpressing it were co-stained for RBMXL3 and FLAG. Confocal microscopy showed strong signal co-localization in cell nucleus (Fig. 1D).

**Figure 1.**
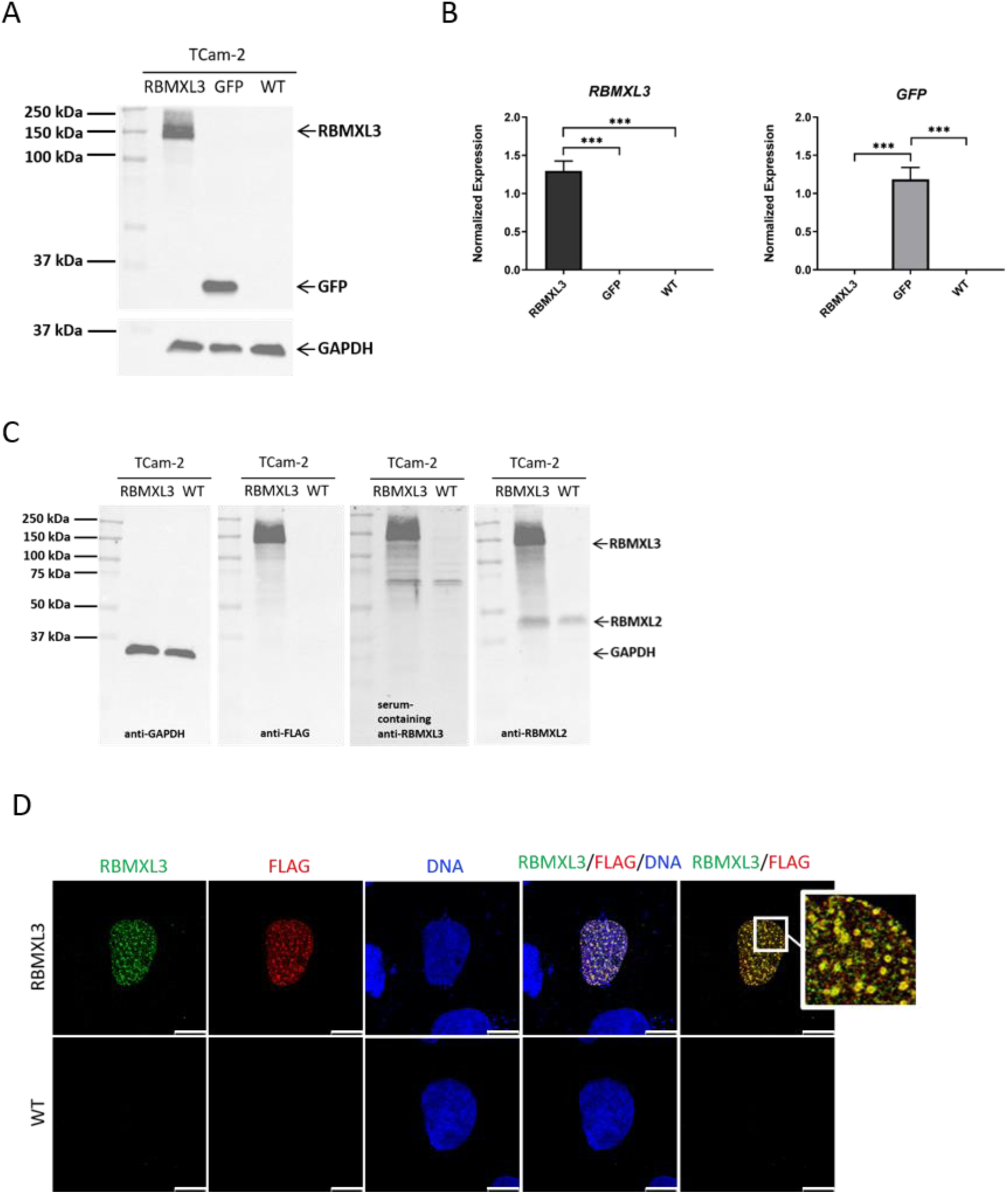
Transient overexpression of *RBMXL3* in TCam-2 cells. (A) Western Blot of transfected (RBMXL3, GFP) and wild-type (WT) TCam-2 cells using anti-FLAG antibody which detects the RBMXL3 and GFP, tagged with a FLAG tag. GAPDH served as a loading control. (B) Expression of *RBMXL3-FLAG* and *GFP-FLAG* in TCam-2 cells on mRNA level, assayed 48h after transfection. Normalized to *GAPDH* and *ACTB*. Ordinary one-way ANOVA non-parametric test with Tukey’s multiple comparisons test was used for statistics. Error bars represent standard deviation (SD) with n = 3, ***p < 0.001. (C) Western Blot of TCam-2 cells detecting RBMXL3 protein with the use of anti-FLAG, anti-RBMXL3, and anti-RBMXL2 antibodies. GAPDH served as a loading control. The additional band appearing between 75 and 50 kDa was derived from serum which was present in one of two batches of RBMXL3 specific antibody used (Supplemental Fig. S1). (D) Co-immunofluorescence staining using anti-FLAG and specific anti-RBMXL3 antibodies in the wild-type (WT) and *RBMXL3*-expressing TCam-2 cells. The zoom window illustrates signal colocalization of RBMXL3 and FLAG. Scale bar 11.6 μm.

To further confirm RBMXL3 subcellular localization, TCam-2 cells overexpressing the protein of interest were subjected to cell fractionation. Visible enrichment of FLAG-tagged RBMXL3 was detected in the chromatin fraction (Fig. 2A). This result encouraged us to investigate whether RBMXL3 binds DNA by performing chromatin immunoprecipitation sequencing (ChIP-seq). However, the FLAG ChIP-seq did not detect reproducible DNA enrichment relative to the GFP control (unpublished data). This outcome may be influenced by technical constraints, as well as the possibility that RBMXL3 associates with chromatin indirectly or transiently, arguing against stable direct DNA binding.

**Figure 2.**
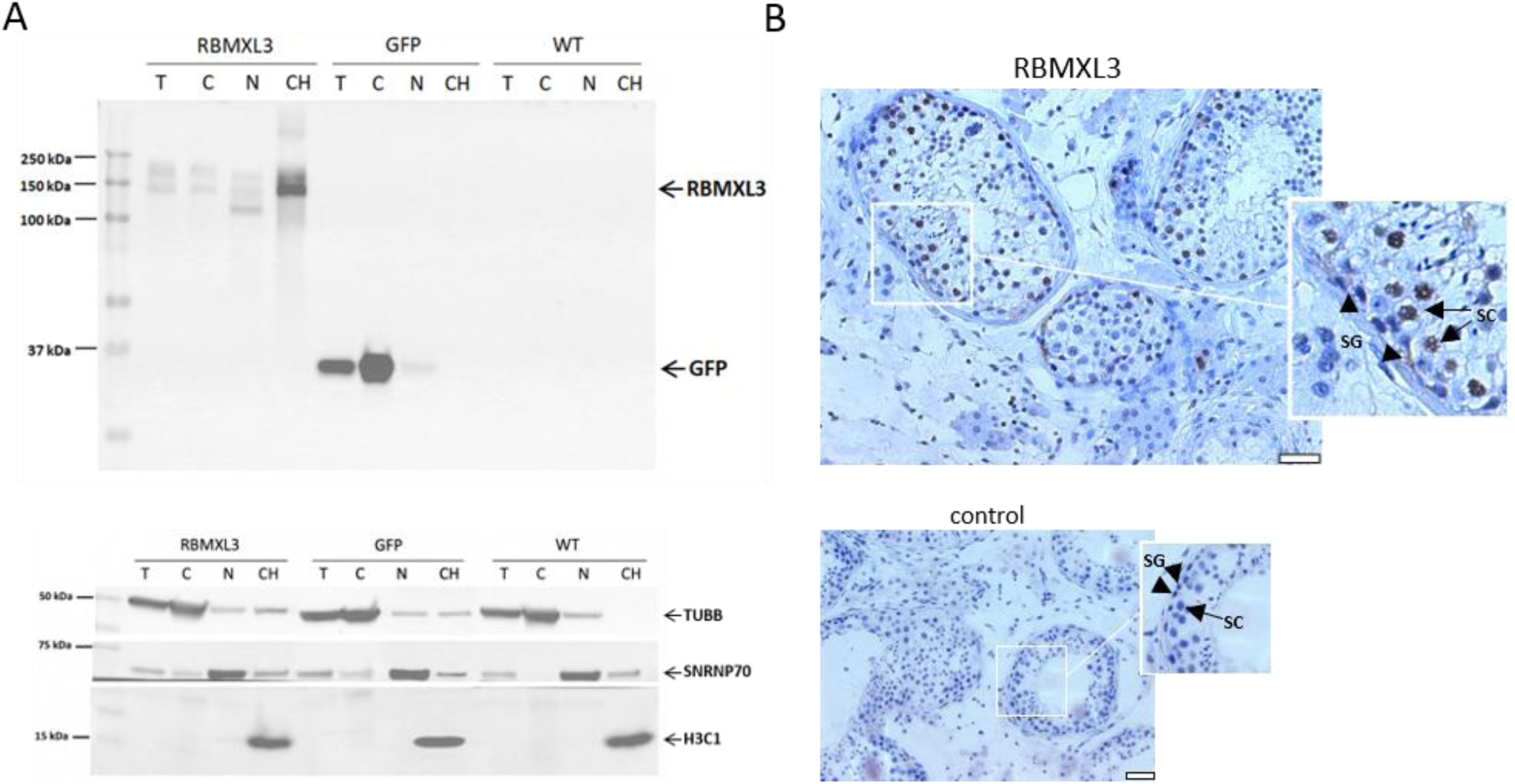
RBMXL3 enrichment in the chromatin fraction of TCam-2 cells. (A) Western Blot analysis using anti-FLAG antibody performed on total (T), cytoplasmic (C), nuclear (N), and chromatin (CH) protein fractions obtained from TCam-2 cells (overexpressing RBMXL3 and two controls: GFP and WT). The subcellular fractions were evaluated using the following markers: tubulin beta class I (TUBB) as a cytoplasmic marker, small nuclear ribonucleoprotein U1 subunit 70 (SNRNP70) as a nuclear marker, and H3 clustered histone 1 (H3C1) as a chromatin marker. In each fraction, there is visible enrichment of the respective marker. (B) Immunohistochemistry of RBMXL3 using specific RBMXL3 antibody in the normal adult human testis. Negative control is presented below. Scale bar 50 μm. SG, spermatogonia (arrowheads); SC, spermatocytes (arrows).

Finally, to confront our observations in TCam-2 cells with *in vivo* RBMXL3 distribution in human testis we performed immunohistochemical staining for RBMXL3 using tissue section of a healthy adult human testis (Fig. 2B). The result was in line with transcriptomic data from the Human Protein Atlas according to which the highest *RBMXL3* mRNA level is detected in spermatogonia and spermatocytes. It is also worth noting that the localization of RBMXL3 protein, although nuclear for spermatocytes, seems to be rather cytoplasmic in the case of spermatogonia. Importantly, previous data from the HPA did not provide exact RBMXL3 protein staining due to the use of an RBMXL2 antibody (with positive signal in spermatogonia, spermatocytes, and spermatids), as the antibody against RBMXL3 is not yet commercially available. To our knowledge, this is the first demonstration of immunohistochemistry-based localization of RBMXL3 using an RBMXL3-specific antibody. In summary, the presence of RBMXL3 in the human testis supports its potential role in the spermatogenesis process, during which precise alternative splicing regulation and genome integrity maintenance are crucial and remain the major challenges.

### RBMXL3 influences gene expression, mRNA splicing, and interacts with protein factors involved in RNA metabolism

To investigate how RBMXL3 protein affects the human transcriptome we first performed RNA-sequencing using RNA from TCam-2 cells expressing either *RBMXL3* or *GFP* (both modified with lentivirus) as well as from WT cells (Fig. 3A,B; Supplemental Table S1; Supplemental Fig. S2A). Since both the GFP and WT TCam-2 cells were used as controls we focused on genes that were the differentially expressed genes (DEGs) in both comparisons i.e. ‘RBMXL3 vs. GFP’ and ‘RBMXL3 vs. WT’ (see Materials and Methods section for more details). Using the cutoff |log2FC| ≥ 1 and FDR < 0.05 we identified 96 up-regulated and 74 down-regulated genes after *RBMXL3* overexpression (Fig. 3C; Supplemental Table S2, Supplemental Fig. S2B,C). The transcriptome analysis detected both protein-coding and long non-coding RNA (lncRNA) genes to be differentially expressed (Fig. 3D). Among DEGs there were transcripts coding for proteins that contribute to the structural integrity of the extracellular matrix (e.g. COL3A1, COL7A1, COL9A3, FBN2, LUM, THBS1, TNXB) which might be important in the context of differentiating germ cell migration during spermatogenesis. Interestingly, after lentiviral transduction we observed differences in the RBMXL3 band pattern on WB compared with that observed after transient transfection (Fig. 1A and 3A). Although the underlying mechanism is difficult to determine, this effect may result from enhanced protein degradation in the stable cell line. The lower-molecular-weight bands likely represent degradation products and are detectable only in cells with long-term *RBMXL3* expression and prolonged RBMXL3 activity.

**Figure 3.**
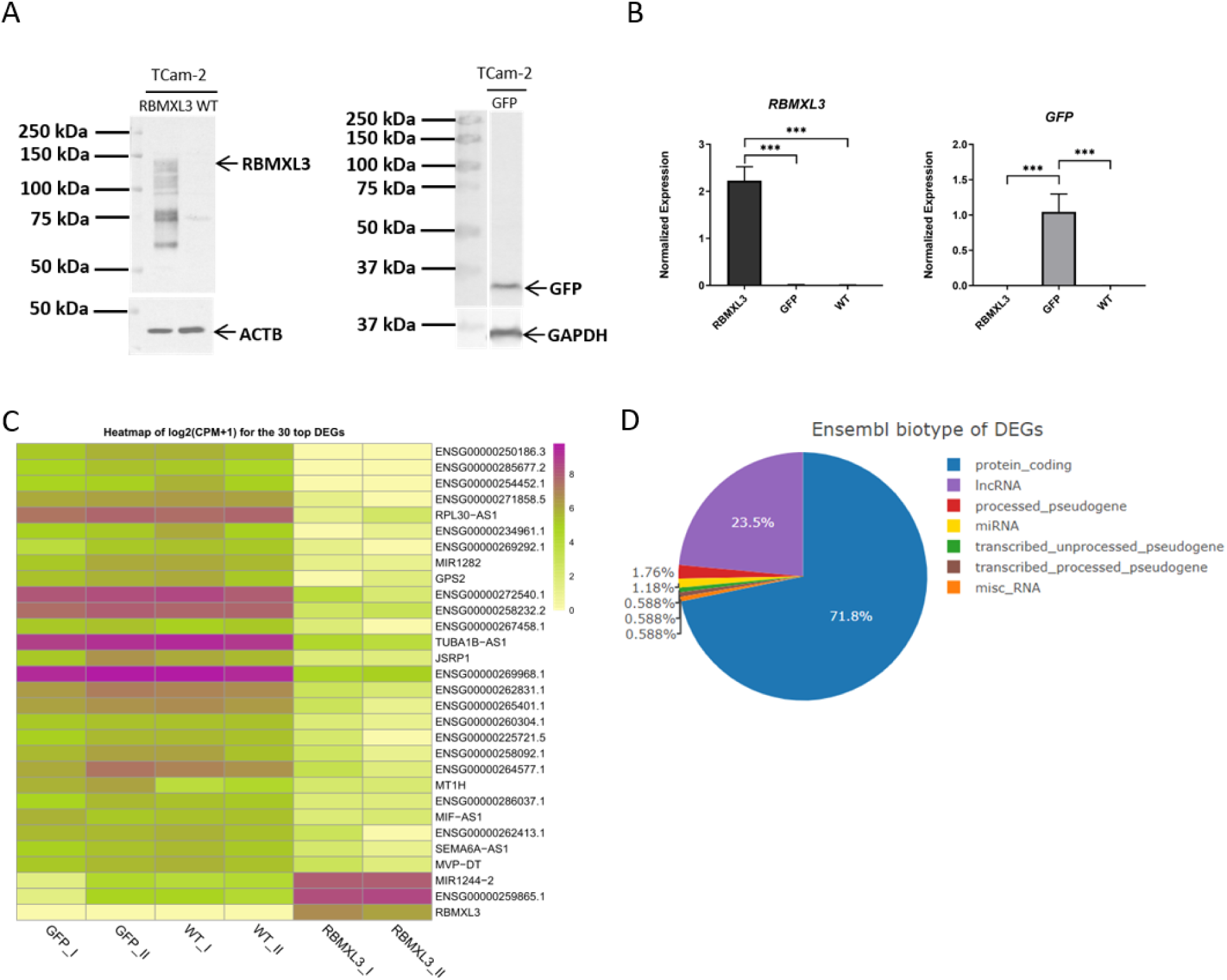
RBMXL3 drives transcriptome changes in human cells. (A) Western Blot of TCam-2 cells with stable expression of *RBMXL3-FLAG* and *GFP-FLAG* as well as wild type (WT) using anti-FLAG antibody. GAPDH and ACTB were used as loading controls. (B) The expression level of *RBMXL3-FLAG* and *GFP-FLAG* in stable TCam-2 cell lines investigated at RNA level by RT-qPCR. Normalization was done to *GAPDH* and *ACTB*. Ordinary one-way ANOVA non-parametric test with Tukey’s multiple comparisons test was used for statistics. Error bars represent standard deviation (SD) with n = 3, ***p < 0.001. (C) Heatmap of log2(CPM+1) for the top 30 differentially expressed genes (DEGs) based on RNA-seq analysis of TCam-2 cells, either WT or overexpressing *RBMXL3-FLAG* or *GFP-FLAG* (|log2FC| ≥ 1|, FDR < 0.05). The top 30 DEGs presented on the heatmap were selected based on the highest |log2FC| and the lowest FDR values derived from the ‘RBMXL3 vs. GFP’ comparison (Supplemental Table S1,2). (D) The Ensembl biotype of DEGs based on RNA-seq analysis of the TCam-2 cells.

Given the spermatogenesis-specific role of RBMXL2 in splicing regulation in mice [13] and the sequence similarity between human RBMXL2 and RBMXL3 (85% and 56% at the DNA and protein levels, respectively), we tested whether RBMXL3 can also play a role in this process. We analyzed our RNA-seq data using SUPPA2 software to identify differential alternative splicing (AS) events. We focused on significant predicted splicing events present in both comparisons (‘RBMXL3 vs. GFP’ and ‘RBMXL3 vs. WT’, see Materials and Methods section for more details) and identified 916 splicing events in 470 transcripts (p≤0.05 and |ΔPSI|≥0.15) upon expression of *RBMXL3-FLAG* (Supplemental Table S3,4). The majority of AS events were alternative first exon (493) and exon skipping (174) (Fig. 4A). Next, to extract the transcripts associated with human male gamete generation we utilized the genes list annotated to the GO term (GO:0048232; male gamete generation) and compared them with differentially spliced genes (DSGs) and DEGs. This approach led us to identify 14 genes: *DLD*, *YY1*, *CCNY*, *ATN1*, *GORASP2*, *GAMT*, *SKIL*, *WDR33*, *SMAD4*, *GJA1*, *CEP57*, *BCL2L1*, *DHX36* and *LRRK2* that potentially underwent AS due to *RBMXL3* expression (Fig. 4B; Supplemental Table S5). Gene ontology (GO) [67, 68] enrichment analysis of DSGs showed that they are associated with terms such as mitotic cell cycle process, chromosome organization, spindle, but also with regulation of RNA splicing, spliceosomal complex, transcription factor binding, ncRNA metabolic process and translation, to name a few (Fig. 4C, for the complete list of GO terms see Supplemental Table S6). To further explore the RBMXL3 function we also performed Reactome pathway analysis of DSGs which underlined terms like Nonsense-Mediated Decay (NMD), mRNA splicing, translation, and M phase, among others (Supplemental Fig. S3, for the complete list of significant Reactome terms see Supplemental Table S6). This data indicates that RBMXL3 might play a role in the alternative splicing of key genes, important for spermatogenesis and beyond.

**Figure 4.**
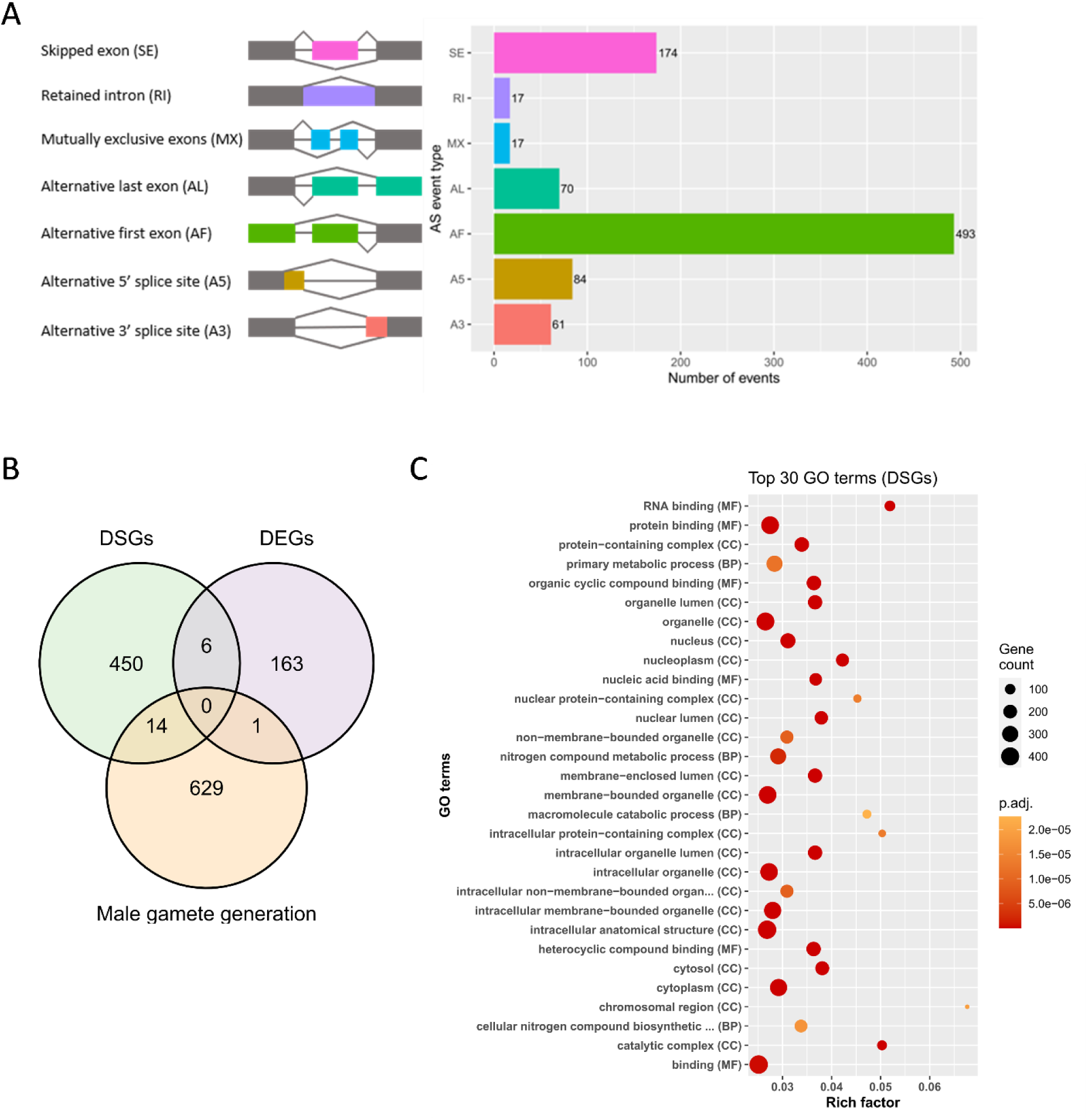
Characterization of differentially spliced genes (DSGs) and alternative splicing (AS) events upon *RBMXL3* expression. (A) SUPPA2 analysis of RNA-seq data from TCam-2 cells identified 916 differential alternative splicing events of different types (p≤0.05 and |dPSI|≥0.15) (Supplemental Table S4). (B) Venn diagram illustrating the total number of DSGs and DEGs based on RNA-seq data in comparison to the list of genes associated with human male gamete generation (Supplemental Table S5). The male gamete generation genes list annotated to GO:0048232 was retrieved from AMIGO2 (accession date 2024-02-02)[115]. Intersection was prepared using Ensembl IDs. (C) Gene ontology (GO) enrichment analysis of differentially spliced genes (DSGs) due to overexpression of the RBMXL3 protein in TCam-2 cells (adjusted p-value < 0.05). For the complete list of GO terms see Supplemental Table S6. The abbreviations in the brackets represent the three GO categories: biological process (BP), molecular function (MF), and cellular component (CC). The rich factor represents the ratio of the number of genes from the input list associated with a given GO term (significant) to the total number of genes annotated to that term (annotated).

To deepen our understanding of the molecular protein interaction network of RBMXL3 we performed RBMXL3-FLAG co-immunoprecipitation followed by mass spectrometry analysis (Co-IP-MS) to identify RBMXL3-interacting proteins (Fig. 5A). This approach allowed us to select 16 potential interactors (Supplemental Table S7). Among them were known splicing factors (SFs) such as EIF4A3, a core component of the splicing-dependent multiprotein exon junction complex (EJC) and translation initiation factor and DDX39B, a protein involved in the processing and nuclear export of mRNA. Some of the potential interactors were also linked to processes such as rRNA biogenesis (e.g. NOLC1) and protein degradation (e.g. PSMD2). Furthermore, analysis performed with the STRING database indicated that most proteins identified by Co-IP-MS create a functional network involved in the regulation of gene expression (Fig. 5B; Supplemental Table S8). In addition, GO annotation performed on the Co-IP-MS detected proteins highlighted terms such as: negative regulation of RNA binding, U4 snRNP, U2-type prespliceosome assembly, nuclear-transcribed mRNA catabolic process, nonsense-mediated decay, translation release factor activity, codon specific and cytoplasmic translational termination (Fig. 5C; Supplemental Table S9). Moreover, according to the original STRING database network for RBMXL3, the majority of predicted interactors are small nuclear ribonucleoproteins (snRNPs), which are core components of the spliceosome (Supplemental Fig. S4A), further supporting the involvement of RBMXL3 in RNA splicing. However, it is worth noting that 7 out of 10 predicted interactors are shared between RBMXL3 and RBMXL2, which may arise also from sequence similarity between these proteins.

**Figure 5.**
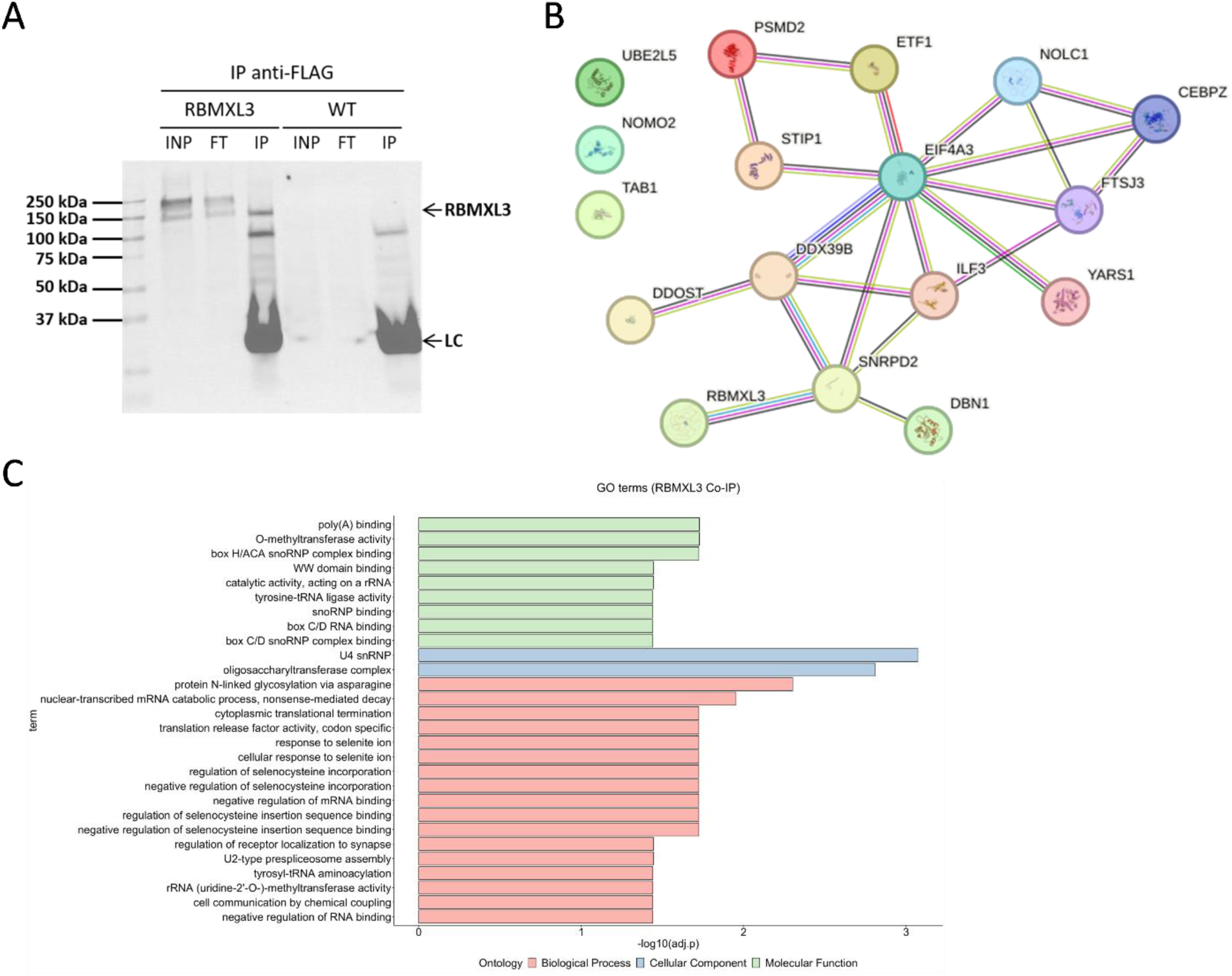
Identification of RBMXL3 protein interactors by Co-IP-MS. (A) Representative Western blot after immunoprecipitation using magnetic beads coated with anti-FLAG M2 antibody. RBMXL3-FLAG in the input (INP), flow-through (FT), and immunoprecipitated (IP) fractions was detected using FLAG antibody. LC - light chains of antibody. (B) Protein-protein interaction (PPI) network created among 16 identified potential protein interactors and RBMXL3 based on the STRING database (Supplemental Table S7,8). The 0.4 cutoff score was used to filter out interactions. (C) GO analysis of proteins identified by Co-IP-MS. GO terms with adjusted p-value ≤ 0.05 were considered significant (Supplemental Table S9). The colors represents the three GO categories (green: MF, blue: CC, red: BP).

For further validation, two interactors (involved in splicing) were chosen. We used immunofluorescence co-staining and confocal imaging of the two selected proteins, EIF4A3 and DDX39B also known as UAP56 (Supplemental Fig. S4B,C,D). Although signal co-localization between RBMXL3 and both interactors was not ubiquitous, we observed several co-localization events. It is worth noting that two DDX39B antibodies (both polyclonal, coming from different manufacturers) had different molecular patterns of staining.

### Identification of RBMXL3 targets by eCLIP

In order to identify RNAs directly interacting with RBMXL3 we performed enhanced crosslinking and immunoprecipitation (eCLIP) followed by high-throughput sequencing [54]. Using FLAG antibody, two biological replicates of each sample, TCam-2 cells transfected with *RBMXL3-FLAG* or *GFP-FLAG* (as control) were subjected to eCLIP (Fig. 6A). For comprehensive analysis, we focused on binding sites that intersected between the two ‘RBMXL3’ replicates and were not detected in any of the ‘GFP’ replicates. We identified more than 1,200 annotated transcripts with over 6,900 RBMXL3-binding peaks (Supplemental Table S10,11). As the RBMXL3-binding site position may be different within the same transcript (e.g. *MSH3*) between the two replicates and thus do not meet the positional intersection condition we prepared an extended list of transcripts intersected not only at the positional level but also at the transcripts level (Supplemental Table S11). Distribution of enriched peaks in the RBMXL3-FLAG IP sample showed that the majority of RBMXL3 binding sites are present within introns, suggesting the role of RBMXL3 in splicing regulation (Fig. 6B). Moreover, eCLIP data suggest that RBMXL3 interacts with both protein-coding and long non-coding RNAs (lncRNAs) (Fig. 6C). Enriched motif analysis revealed that the top two most enriched represent the 5’ and 3’ splicing signal consensus (Fig. 6D). We have also analyzed the profile of eCLIP peaks binding to the vicinity of the RNA splice sites and show that there is a visible enrichment near splice sites, further supporting the role of RBMXL3 in splicing (Fig. 6E). Analysis of eCLIP peaks coverage on an mRNA model showed a strong enrichment of RBMXL3 binding observed also around the 5’UTR region and start codon (Supplemental Fig. S5A). Then, we again employed the male gamete generation genes list by comparing it with transcripts bound by RBMXL3 and identified 32 genes, including *BRCA2, CLOCK, MSH6, TSGA10, ATRX, MLH3, KNL1, SETX, JAM3, NECTIN2, TCFL5, IGF2R, TDRD5,* and *CDYL*, among others (Fig. 6F, Supplemental Table S12). Moreover, we investigated how many transcripts of the differentially spliced genes (DSGs) were directly bound by RBMXL3 based on eCLIP data and identified 48 transcripts that might be direct targets of RBMXL3 protein, including*HSPH1, CENPJ, HDAC2, FTX, TFDP2, PLEKHA5, CBFB, RIF1, E2F5, ZNF131, SP4,* and *CEP57*, among many others (Fig. 6G; Supplemental Table S13). In this group of genes, only one (CEP57) was assigned to the male gamete generation GO term. However, it is important to emphasize that many genes important for this process in humans still remain to be discovered and properly annotated. Functional characterization of identified RNAs using GO revealed that targets are involved in processes such as sister chromatid segregation, nuclear chromosome segregation, spindle organization, mitotic cell cycle process, and stem cell population maintenance (Fig. 6H, for the complete list of GO terms see Supplemental Table S14). These data indicate that RBMXL3 may play a crucial role in human spermatogenesis, particularly in germ cell differentiation, regulating the processes important for stem cells biology. Other significantly enriched GO terms among RBMXL3 RNA targets included the transcription regulator complex, regulation of RNA splicing, DNA-binding transcription factor activity, protein methylation, regulation of protein deacetylation and epigenetic regulation of gene expression. To further understand how the functions of RBMXL3-bound RNAs may relate to spermatogenesis we performed Reactome pathway analysis. This identified terms like centrosome maturation, separation of sister chromatids, and recruitment of mitotic centrosome proteins and complexes, pathways which are crucial to ensure gamete quality (Supplemental Fig. S5B, for the complete list of significant Reactome pathways see Supplemental Table S14). It is worth noting that among the detected RBMXL3 targets we found the RBMXL3 mRNA, suggesting that the RBMXL3 binds its own transcript, as several RNA-binding proteins (RBPs) do. Summarizing, by using eCLIP-seq we confirmed that RBMXL3 is a functional RBP and identified its RNA targets. Our data indicate that the primate-specific RBMXL3 protein exerts a significant impact on human cell transcriptome.

**Figure 6.**
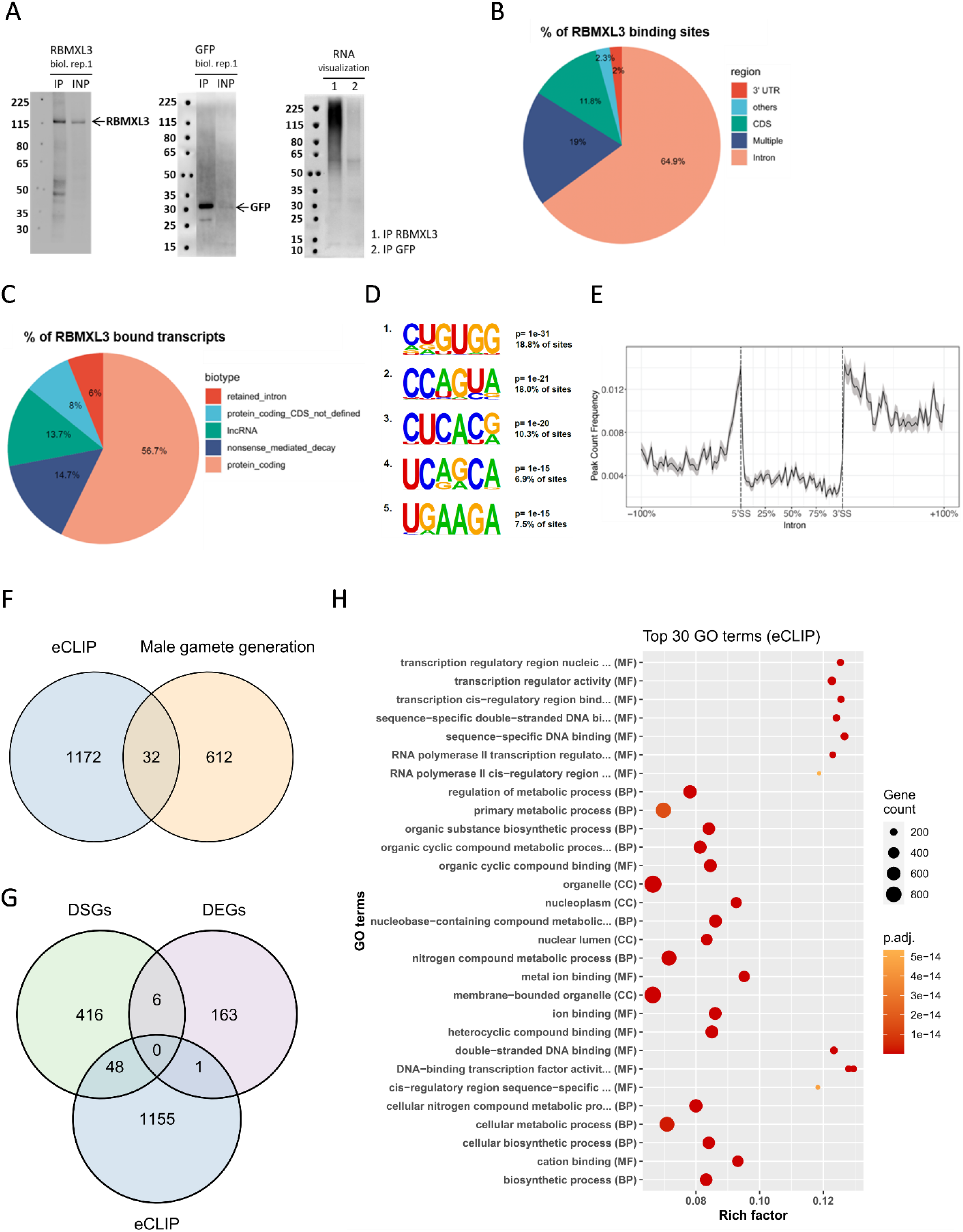
Global profiling of RNA target sites directly interacting with RBMXL3 by eCLIP. (A) Immunoblotting of RBMXL3-FLAG using anti-FLAG antibody (left) and visualization of immunoprecipitated RNA (right) in TCam-2 cells. (B) Distribution of the RBMXL3 binding sites in the different regions. All eCLIP peaks that map to a specific genomic location with multiple, different transcripts (thus possessing the same or different biotypes) are within the “Multiple” category. (C) The Ensembl biotype of the RBMXL3-bound transcripts in TCam-2 cells. (D) Significant binding motifs found in at least 5% of binding sites, sorted by p-value and identified by HOMER *de novo* motif analysis. (E) Visualization of the RBMXL3 RNA-binding sites with eCLIP peaks coverage in the vicinity of the RNA splice sites. (F) Venn diagram of transcripts bound by RBMXL3 and genes associated with human male gamete generation. The male gamete generation genes list annotated to GO:0048232 was retrieved from AMIGO2 (accession date 2024-02-02) [115] (Supplemental Table S12). Intersection was prepared using Ensembl IDs. (G) Venn diagram illustrating the total number of DSGs and DEGs based on RNA-seq data in comparison to the list of transcripts detected in RBMXL3 eCLIP thus directly bound by RBMXL3 (Supplemental Table S13). Intersection was prepared using Ensembl IDs. (H) GO analysis for the transcripts bound by RBMXL3 based on eCLIP data. For the complete list of significant GO terms see Supplemental Table S14 (adjusted p-value < 0.05). The rich factor represents the ratio of the number of genes from the input list associated with a given GO term (significant) to the total number of genes annotated to that term (annotated).

### The primate-specific RBMXL3 protein suppresses human LINE-1 retrotransposition

We analyzed the eCLIP data in regards to repetitive elements and showed that RBMXL3 binds sequences belonging to the L1 (Long Interspersed Nuclear Element-1 also known as LINE-1) family of retrotransposons (Fig. 7A). This result encouraged us to investigate the effect of RBMXL3 on L1 retrotransposition by performing a plasmid-based L1 retrotransposition assay using HEK293T cells (see Materials and Methods section for more details). We observed that the RBMXL3 protein heavily reduces L1 retrotransposition (Fig. 7B) [66] in this assay, indicating that it may play a protective role against LINE-1 mobilization, which is especially important in the context of germline and spermatogenesis [69]. To confirm that this strong effect of RBMXL3 on L1 retrotransposition is a true positive result we performed a series of control experiments. The lack of statistically significant changes in the percentage of transfected cells indicated that RBMXL3 plasmid and expression do not significantly affect the transfection process or generally reduce EGFP and mCherry protein production (Supplemental Fig. S6A,B). In fact, we detected higher EGFP and mCherry intensities in the RBMXL3 expression condition which currently cannot be explained. To complement this result, we also tested whether RBMXL3 affects endogenous L1 mRNA levels. Based on our RNA-seq data, we analyzed L1s expression in TCam-2 cells expressing *RBMXL3*, using *GFP*-expressing and WT cells as controls (Fig. 7C; Supplemental Table S15). Our analysis showed that some L1s are differentially expressed in sample with RBMXL3, but no subfamily displayed a similar up- or down-regulation pattern between the two comparisons when using a threshold of |log2FC| ≥ 0.585, FDR ≤ 0.05. We did not observe significant changes in the expression of endogenous L1HS class (also called L1PA1 and Ta) [70–72] of retrotransposition competent L1s in humans (of which the L1RP element used in the plasmid-based assay is a representative). Overall, our data provide evidence that RBMXL3 participates in the effective silencing of transposable elements (TEs), specifically L1, which is important for safeguarding spermatogenesis.

**Figure 7.**
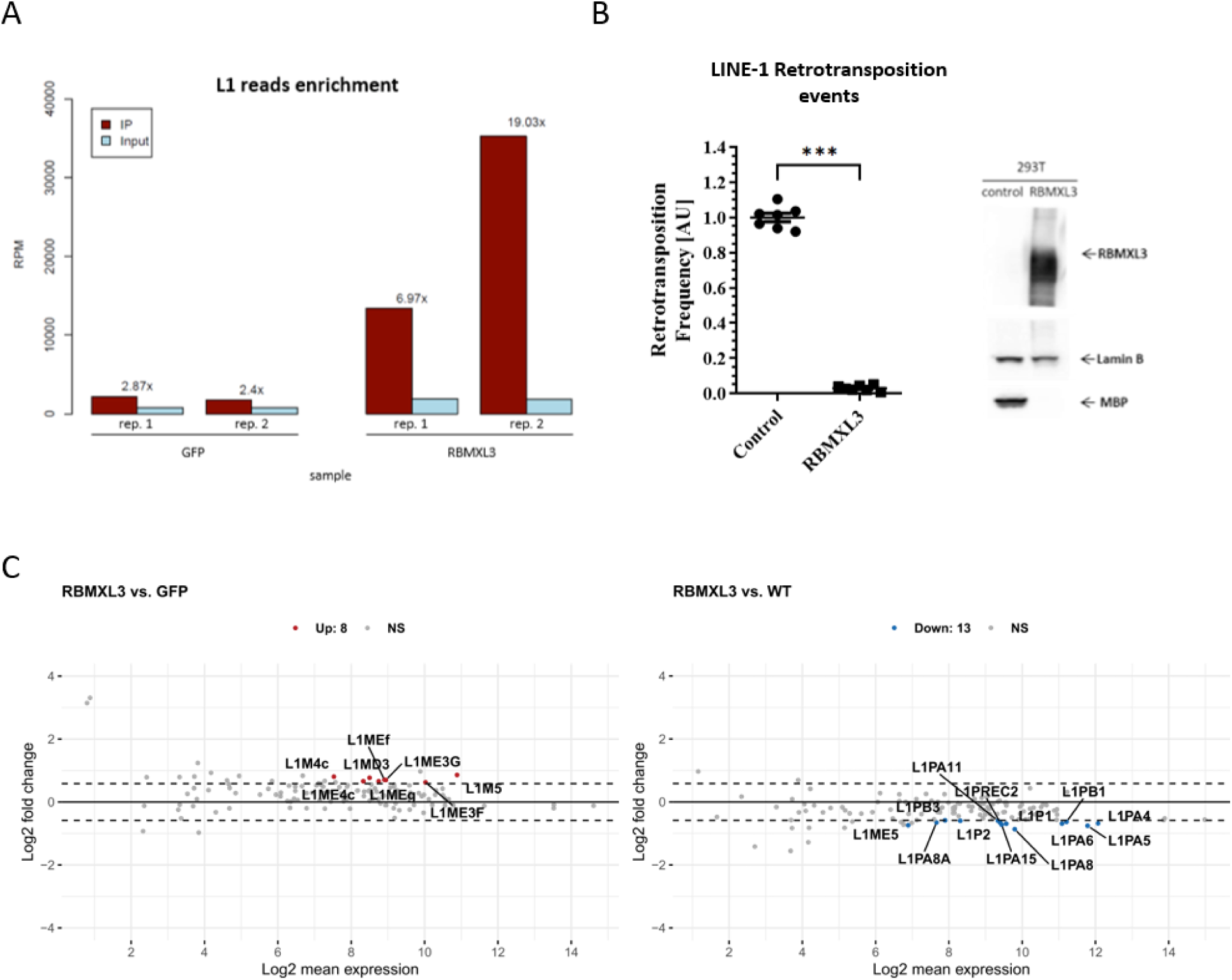
RBMXL3 binds sequences belonging to L1 family and heavily reduces L1 retrotransposition. (A) L1 reads enrichment in IP vs. Input sample based on repetitive elements analysis of the eCLIP data indicates that RBMXL3 protein binds sequences belonging to the L1 family. (B) Effect of overexpression of *RBMXL3* and *MBP* (as control) on L1 retrotransposition in HEK293T cells using L1 retrotransposition reporter assay. The results of independent experiments were normalized relative to the control (MBP). *RBMXL3* expression induces a ∼15-20 fold reduction in L1 retrotransposition; each point represents a biological replicate, ***p<0.001 (left). Western blot analysis using anti-FLAG, anti-Lamin B and anti-MBP antibodies to detect respectively: RBMXL3, Lamin B (loading control), and MBP protein (right). (C) MA plots showing the L1s from RNA-seq analysis using TEtranscripts package (Supplemental Table S15). Differentially up-regulated L1s (|FC| ≥ 1.5, FDR ≤ 0.05) are labeled red, whereas differentially down-regulated L1 (|FC| ≤ - 1.5, FDR ≤ 0.05) are labeled blue. Horizontal dashed lines indicate log2(1.5) or -log2(1.5).

## Discussion

This study presents novel findings on the primate-specific *RBMXL3* and its critical role in RNA metabolism. We confirmed the nuclear localization of RBMXL3 protein in TCam-2 cell line and its presence in spermatogonia and spermatocytes within the human testis. For the first time, we demonstrate that RBMXL3 acts as an upstream regulator of RNA metabolism by binding to a wide range of RNA molecules in human TCam-2 cells. Our data reveal that RBMXL3 drives transcriptomic changes by influencing both gene expression as well as alternative splicing of multiple transcripts. By using LINE-1 retrotransposition reporter assays we discovered that RBMXL3 suppresses L1 retrotransposition in human cells. Although the TCam-2 cell line serves as an important *in vitro* model of human germ cell biology [42, 73], it does not recapitulate the complex differentiation program of germ cells within the human testis. Therefore, we interpreted the TCam-2 data primarily as indicative in the context of human spermatogenesis and focused on gene/protein targets that are expected to be present in germ cells where RBMXL3 is expressed *in vivo*.

Alternative splicing regulation has been shown to be pivotal in the context of differentiation processes [74] and is crucial for proper spermatogenesis [13, 42, 75–77]. We and others identified mutations in *RBMXL3* gene found in non-obstructive azoospermia (NOA) patients with Sertoli cell-only syndrome (SCOS) [5] and recently in patients with asthenozoospermia. By using modified TCam-2 cells as a model of human primordial germ cells we demonstrate the role of RBMXL3 in regulation of transcriptome. Our analyses led to identification of potential RBMXL3 target transcripts some of which might undergo AS depending on RBMXL3 presence. Among potential targets is *FOS* (also known as *c-FOS*), a protooncogene involved in cell proliferation [78], which was highly enhanced upon *RBMXL3* overexpression. FOS protein is present in the human testis and has been linked to spermatogenesis [79]. It is decreased in the testis of men with NOA. Furthermore, among the genes detected as potentially differentially spliced in TCam-2 cells upon RBMXL3 overexpression, we found the *YY1* transcription factor. This gene has been shown to influence the heterochromatin in mouse spermatocytes which affects genome stability during male meiosis [80]. It has been also suggested that YY1 is a key factor for maintaining the viability of spermatogonia and the stemness of spermatogonial stem cells (SSCs) [81]. In patients with *RBMXL3* sequence defects, the precise control of YY1 during spermatogenesis might be disrupted, which in theory could lead to exhaustion of the spermatogonia pool and azoospermia. Another interesting gene found in the group of differentially spliced genes (DSGs) was *GJA1*, coding for the predominant testicular gap junction protein involved in the regulation of fate decisions in human SSCs [82]. Our data show also that RBMXL3 may influence AS of *HNRNPH1*, a known splicing factor. In mice, HNRNPH1 has been shown to be involved in the establishment of crosstalk between Sertoli cells (SCs) and germ cells, further, conditional knock-out of this gene in SCs resulted in mice infertility [83]. Another study confirmed the presence of HNRNPH1 protein in the human testis, detecting it mainly in the nucleus of spermatocytes and indicating its role in the suppression of spermatogenesis induced by heat exposure [84]. Other examples of RNA-binding proteins involved in spermatogenesis are RBMXL2 [13] and RBM46 [85], whose knockout causes male infertility in mice. RBMXL2 has been implicated in suppressing aberrant splice sites during mouse spermatogenesis to ensure proper sperm production [13]. Our data suggest that RBMXL3 might play an important role in spermatogenesis and that the variants in the *RBMXL3* gene may potentially disturb normal protein function, also influencing the pool of RBMXL3-regulated RNAs and lead to male infertility.

Studies concerning human spermatogenesis are extremely limited due to the lack of comprehensive *in vitro* models of spermatogenesis and the fact that many genes crucial for this process still remain to be discovered. Although the functions of the majority of specific isoforms are not known yet, precise regulation of gene expression and AS is necessary to ensure the successful completion of male gamete formation. Taking into consideration that SCOS (diagnosed by testicular biopsy in men with *RBMXL3* variants) is generally characterized by a lack of germ cells in the testis, based on our data, we conclude that RBMXL3 may play a role in processes important for spermatogonial stem cells (SSCs) differentiation towards spermatozoa and/or their self-renewal. In patients with disrupted RBMXL3 function, these processes might be affected, leading to spermatogenic impairment.

In our eCLIP data we identified more than 1,200 transcripts with over 6,900 RBMXL3 binding sites. Among the RBMXL3 RNA targets, we found multiple transcripts coding for proteins involved in DNA repair pathways. Two of these, important in the context of spermatogenesis, are BRCA1 and BRCA2, which have established roles in cancer but are also involved in meiotic sex chromosome inactivation during spermatogenesis and meiotic homologous recombination [86, 87]. Another interesting examples could be the *MSH2* and *MSH6* transcripts, coding for mismatch repair proteins and also expressed during spermatogenesis. These proteins ensure the correction of replication errors predisposing to different malignancies and participate in the genomic integrity and stability maintenance in male germline [88]. Among the transcripts bound by RBMXL3 we also found the *EP300* mRNA (bound within its coding sequence). The protein product of this gene functions as a histone acetyltransferase that regulates transcription via chromatin remodeling and is important in the context of human stem cells and differentiation [89, 90]. Moreover, the knockdown of *EP300* reduced stem cell content in primary cultures of undifferentiated spermatogonia from mice [91]. Another RBMXL3 RNA target is *PREP*, bound within its intronic sequence. PREP is recognized as a multifunctional protein, which has been detected in human testicular tissue [92] and numerous reports suggested its role in mouse, rat, and also human spermatogenesis [92, 93]. Another potentially interesting RNA target in the context of NOA and SCOS might be *FANCM*, a testis-enhanced gene that has already been related to such phenotype in human male patients [94]. Our data indicate that RBMXL3 protein participates in upstream regulation of many key genes important for spermatogenesis, where precise regulation of sophisticated gene expression patterns is necessary to produce haploid spermatozoa. In the wide pool of RNAs bound by RBMXL3 (eCLIP data) we also found *FOXJ3* (a transcription factor expressed in human testis). Studies of *Foxj3* in mice have shown that deletion of *Foxj3* from spermatogonia caused SCOS in males and deletion from spermatocytes led to meiotic arrest (both models exhibited complete sterility) [95]. The conclusion was that FOXJ3 is required for spermatogonia survival and participates in meiosis. According to our eCLIP data, RBMXL3 binds *FOXJ3* RNA within its intronic sequence, which may influence its post-transcriptional regulation as well as downstream modifications, specific for humans. In NOA patients with RBMXL3 impaired function such disruption potentially may lead to SCOS phenotype. Any DNA sequence variants found in RBMXL3 may potentially affect the wide repertoire of RBMXL3 targets and perturb the human male germ cell differentiation process, which still remains to be fully elucidated. Our eCLIP data also suggest that RBMXL3 may contribute to the epigenetic regulation of gene expression as it binds a wide range of RNAs coding for proteins involved in the regulation of histone modifications such as PRDM2, PRMT2, CREBBP, TET1, and TET2. This is important in the context of the spermatogenesis process which is accompanied by alterations in histone modifications in order to modulate gene expression changes during gamete differentiation [96, 97] and might be disrupted in patients with *RBMXL3* mutations. The RBMXL3 interplay with epigenetic pathways might also contribute to the silencing of transposable elements [98].

Although the overlap between RBMXL3-induced DSGs (RNA-seq) and RBMXL3-bound transcripts (eCLIP) is relatively low, we found proof for direct binding of RBMXL3 to almost 10% of potential DSGs. RBMXL3 may participate in splicing of these transcripts by interacting with other proteins, for example, known splicing factors (EIF4A3, DDX39B) that we identified in our Co-IP-MS experiment. Moreover, eCLIP peak coverage showed an enrichment of RBMXL3 binding around the 5’ UTR regions and the start codon. This observation may be particularly relevant given the high abundance of alternative first exon (AF) events in our analysis. Such events are common in RNA-seq studies, particularly in complex transcriptomes, and often reflect alternative transcription start site and promoter usage [99, 100]. However, as RNA-seq results capture steady-state transcript levels and are influenced by post-transcriptional regulation, the underlying cause cannot be conclusively determined. Considering the protein-protein interaction of RBMXL3 with translation factor EIF4A3, it is also possible that RBMXL3 may modulate translation in primates. Additionally, our RNA-seq and eCLIP-seq data indicate that RBMXL3 affects not only protein-coding transcripts but also lncRNAs and although their role in spermatogenesis remains largely unknown [101] it is currently certain that they are important players in regulation of many crucial cellular processes, stem cell maintenance and differentiation [102]. Finally, we acknowledge the small sample size used in this study (two biological replicates per condition), which may limit the interpretation of the sequencing data.

Furthermore, a recent study examined the genomic profiles of *MYCN* non-amplified neuroblastoma and revealed novel recurrently mutated genes, and among them was *RBMXL3*, mutated in 19% of samples [35]. As neuroblastoma is the most common extracranial solid tumour in infants and children [103], it is possible that the *RBMXL3* mutations may affect cellular differentiation programs. This could relate to the RBMXL3 protein binding to *ZFHX3* mRNA, coding for a transcription factor (known also as a neuronal differentiation marker) which has been shown to affect different types of cancers. *ZFHX3* might have an opposing influence being associated with a suppressor or oncogenic activity, for example in prostate and breast cancer separately [104]. *ZFHX3* has been also shown to affect the expression of some pluripotency factors and stemness markers, including *MYC*, *TBX3*, *OCT4*, *NANOG*, and *SOX2* [104].

Finally, by using a well-established retrotransposition reporter assay, we show that RBMXL3 expression heavily reduces L1 retrotransposition in human cells. Since during spermatogenesis epigenetic reprogramming occurs leading to erasure of epigenetic silencing marks [98, 105] L1s might regain their transcriptional activities. This suggests that RBMXL3 expressed specifically in spermatocytes and spermatogonia at the time of L1 transcriptional activation might contribute to a yet unknown L1 regulatory mechanism. This further supports the concept of RBMXL3 importance in maintaining genome integrity. Although the mechanism of this action remains unknown, we propose here a few possible scenarios. Firstly, we identified significant enrichment of L1 with RBMXL3 in our eCLIP data, demonstrating direct interaction between RBMXL3 and L1 transcripts and implying a post-transcriptional regulatory mechanism [106]. Although we did not identify L1 retrotransposon proteins in our proteomic data, RBMX has been identified among the proteins associated with the L1 ORF1 protein in human cells [107]. It is therefore possible that RBMXL3 interacts with L1-encoded proteins and/or manages their posttranslational modifications restricting L1 mobilization. A similar mechanism has been shown, for example, for mouse TEX19.1 in pluripotent mouse embryonic stem cells [44]. We also cannot exclude the possibility that RBMXL3 is involved in piwi-interacting RNAs (piRNAs) biogenesis pathways as one of its possible interactors (DDX39B/UAP56) participates in germline piRNAs production and transposon silencing in *Drosophila* [108]. Involvement of RBMXL3 in splicing of *YY1* might affect retrotransposition as YY1 is a known L1 regulatory factor [109]. In conclusion, the postulated RBMXL3 interplay with L1s seems to be a worthwhile direction to follow in the context of germ cell development and beyond.

The discrepancies concerning RBMXL3 detection in transcriptome- and proteome-wide studies indicate the need for a more in-depth analysis of RBMXL3 expression patterns. As RBMXL3 is a primate-specific gene the sense of RBMXL3 research on non-primates animal models is questionable. It is also worth noting that the RBMXL3 protein action might be cell cycle phase-specific, hence the experiments using the phase-arrested human cells would be very beneficial. This might lead to the discovery of new protein interactors and RNA targets. To further investigate RBMXL3 functions more in detail, we would suggest utilizing human stem cell-based approaches and advanced human spermatogenesis models [110, 111] to capture its natural molecular environments and possible transient existence. The extensive repertoire of RBMXL3 RNA targets identified in the eCLIP experiment, spanning a broad range of biological processes, suggests that RBMXL3 expression may be more widespread than currently appreciated, yet remains challenging to detect.

Finally, it is worth noting that RBMXL3 binds its own mRNA as demonstrated by our eCLIP data. Such self-regulatory RBP interaction mode is common among many other RBPs emerging as pivotal post-transcriptional regulators of gene expression [112–114]. In agreement with a possible regulatory feedback loop of RBMXL3 repressing its own mRNA translation we encountered several technical difficulties with maintaining RBMXL3 expression at the protein level, but not at the mRNA level in TCam-2 cells following transduction. We hypothesize that the negative loop might exist between RBMXL3 protein and its RNA which may be the reason why RBMXL3 has such an elusive nature across human tissues and its fine-tuning physiological role.

## Conclusions

Our study provides the first evidence that the primate-specific RBMXL3 protein acts as an upstream regulator of RNA metabolism through extensive RNA-binding activity in human TCam-2 cells. Furthermore, we found that RBMXL3 expression significantly suppresses L1 retrotransposition in human cells, highlighting its importance in maintaining human genome integrity. Additionally, we confirmed the nuclear localization of RBMXL3 in TCam-2 cells (*in vitro* male germline model) upon ectopic expression and, for the first time, demonstrated the presence of the RBMXL3 protein in human testis. Collectively, our data imply that RBMXL3 contributes to the control of transcriptome dynamics in human male germ cells, with its full biological relevance yet to be explored.

## Supporting information

Supplementary Material - RBMXL3

## List of abbreviations

RNA-seq: RNA sequencing
eCLIP: enhanced crosslinking and immunoprecipitation
Co-IP-MS: Co-immunoprecipitation coupled with Mass Spectrometry
LINE-1: L1, Long Interspersed Nuclear Element-1
WES: Whole-exome sequencing
WGS: Whole-Genome sequencing
NOA: Non-obstructive azoospermia
HPA: Human Protein Atlas
ChIP: Chromatin Immunoprecipitation
IHC: Immunohistochemistry
DEGs: differentially expressed genes
AS: alternative splicing
DSGs: differentially spliced genes
GO: Gene ontology
RBPs: RNA-binding proteins
TEs: transposable elements
SCOS: Sertoli cell-only syndrome
SSCs: spermatogonial stem cells

## Declarations

### Ethics approval and consent to participate

Not applicable.

### Consent for publication

The paper has been read and approved by all authors. All authors approved the submission of this paper to Cellular & Molecular Biology Letters for publication. All authors confirmed that neither the manuscript submitted nor any part of it has been published or is being considered for publication elsewhere.

### Availability of data and materials

The datasets generated and/or analyzed during the current study are available in the GEO repository, GSE294696 (RNA-seq) and GSE308226 (eCLIP-seq). All data generated during this study are included in this published article and are available in Zenodo: https://doi.org/10.5281/zenodo.12788249.

## Authors’ contributions

**Conceptualization:** N.R.; **Methodology:** A.J.B., K.T., N.R.; **Investigation:** A.J.B., K.T., M.D., A.M., Z.E.M., Z.W.; **Formal analysis:** E.D., A.J.B., J.Z.-W. (NGS data); E.I., D.C. (MS data); **Resources:** M.K., M.O., A.Y. (selection of gene of interest based on previous studies), M.H.; **Writing – original draft:** A.J.B.; **Writing – review & editing:** Z.W., E.D., K.T., E.I., A.J.B., J.Z.-W., A.N., T.K., A.Y., Z.E.M., N.R.; **Supervision:** N.R.; **Funding acquisition:** N.R., Z.W.

All authors read and approved the manuscript.

## Acknowledgements

The scientific work was created with the assistance of the infrastructure of Department of Cellular, Computational and Integrative Biology (CIBIO, University of Trento) and Poznan Supercomputing and Networking Center (PSNC). We thank Prof J. Alcorn for providing RBMXL3-specific antibody. We thank Marcin Sajek for help with data deposition and Katarzyna Kiwerska for help with IHC experiment. We thank Augustyn Moliński for support during confocal imaging of TCam-2 cells and Barbara Cybulska for help with experiments. This manuscript has been released as a pre-print at bioRxiv (DOI: https://doi.org/10.1101/2025.09.23.677621), available under a CC-BY-NC-ND license.

## Supplemental Tables

Supplemental Table S1 Normalized counts (CPM) based on RNA-seq of TCam-2 cells.

Supplemental Table S2 Differentially expressed genes (DEGs) based on RNA-seq of TCam-2 cells.

Supplemental Table S3 Normalized transcript-level values (TPM) for TCam-2 cells quantified with Salmon from RNA-seq data.

Supplemental Table S4 Differentially spliced genes (DSGs) based on SUPPA2 analysis of RNA-seq data.

Supplemental Table S5 Genes overlap results for DSGs, DEGs, and genes involved in male gamete generation.

Supplemental Table S6 Gene Ontology and pathway enrichment analysis of DSGs detected upon RBMXL3 overexpression.

Supplemental Table S7 Results of RBMXL3 co-immunoprecipitation followed by mass spectrometry.

Supplemental Table S8 RBMXL3 protein-protein interaction network analysis performed with the STRING database.

Supplemental Table S9 Gene Ontology enrichment analysis based on proteins identified in RBMXL3 co-immunoprecipitation experiment.

Supplemental Table S10 Detailed information on detected RBMXL3 eCLIP peaks.

Supplemental Table S11 List of transcripts bound by RBMXL3 identified by eCLIP.

Supplemental Table S12 Genes overlap results for RBMXL3 eCLIP targets and genes involved in male gamete generation.

Supplemental Table S13 Genes overlap results for DSGs, DEGs, and RBMXL3 eCLIP targets.

Supplemental Table S14 Gene Ontology and pathway enrichment analysis of RBMXL3 eCLIP targets. Supplemental Table S15 L1 differential expression in TCam-2 cells based on RNA-seq data.

Supplemental Table S16 List of primers.

