## Supplementary Material - RBMXL3 for "Multi-omic screening identifies RBMXL3 as a primate-specific RNA-binding protein and candidate regulator of RNA metabolism in human spermatogenesis"

[1] Institute of Human Genetics, Polish Academy of Sciences, Poznan, Poland
[2] Department of Gene Expression, Institute of Molecular Biology and Biotechnology, Adam Mickiewicz University, Poznan, Poland
[3] Department of Mathematical and Statistical Methods, Poznan University of Life Sciences, Poznan, Poland
[4] Department of Cellular, Computational and Integrative Biology (CIBIO), University of Trento, Trento, Italy
[5] DNRF Center for Chromosome Stability, University of Copenhagen, DK-2200 Copenhagen, Denmark
[6] Laboratory of Mass Spectrometry, Institute of Biochemistry and Biophysics, Polish Academy of Sciences, Warsaw, Poland
[7] Department of Genetics and Animal Breeding, Faculty of Veterinary Medicine and Animal Sciences, Poznan University of Life Sciences, Poznan, Poland
[8] Department of Urology, Poznan University of Medical Sciences, Poznan, Poland
[9] Department of OBGYN and Reproductive Sciences, Magee-Womens Research Institute, School of Medicine, University of Pittsburgh, Pittsburgh, PA, USA
[10] Institute of Bioorganic Chemistry, Polish Academy of Sciences, Poznan, Poland

A

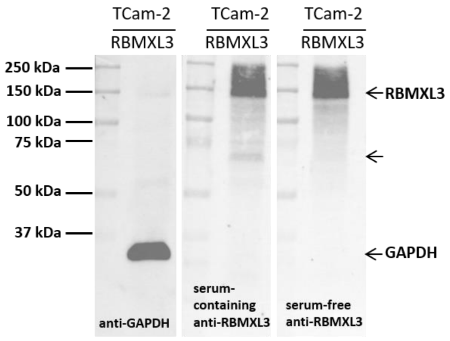


Supplemental Figure S1. RBMXL3 protein detection in TCam-2 cell lysates using RBMXL3-specific antibody. (A) Western Blot of RBMXL3 TCam-2 cells using serum-containing and serum-free RBMXL3 antibody. GAPDH served as a loading control. The additional band appearing between 75 and 50 kDa was derived from serum which was present in one of two batches of RBMXL3 specific antibody used.

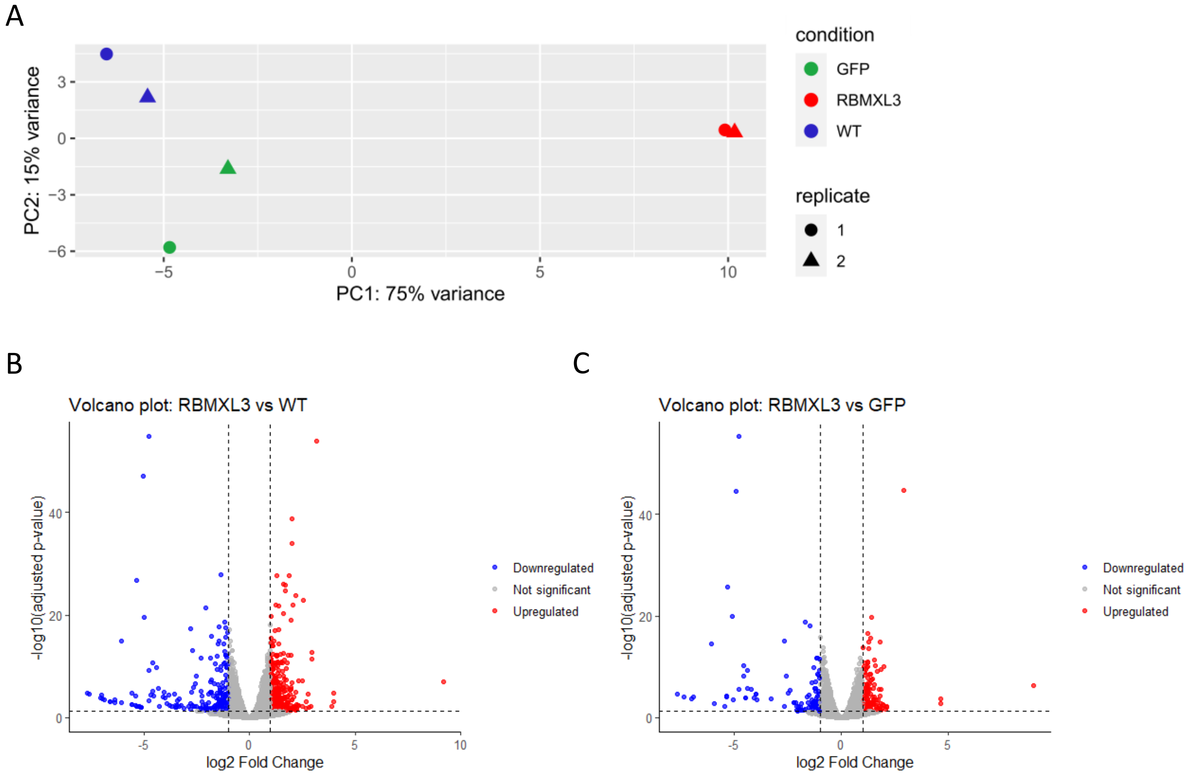


Supplemental Figure S2. Principal component analysis results of TCam-2 RNA-seq data and volcano plots. (A) Principal component analysis (PCA) of the TCam-2 RNA-seq samples. The two biological replicates corresponding to each condition (RBMXL3, GFP and WT) are presented with different shapes. The TCam-2 with RBMXL3 overexpression (highlighted in red) samples separate from the other control samples (GFP and WT, highlighted in green and blue) along principal component 1 (PC1). (B) Volcano plot of differentially expressed genes determined using DESeq2 analysis of RNA-seq data from TCam-2 cells with RBMXL3 overexpression and WT (Supplemental Table S2). Dashed lines represent twofold change in expression (vertical lines) and adjusted FDR < 0.05 cutoff (horizontal line). (C) Volcano plot of differentially expressed genes determined using DESeq2 analysis of RNA-seq data from TCam-2 cells with RBMXL3 and GFP overexpression (Supplemental Table S2). Dashed lines represent twofold change in expression (vertical lines) and adjusted FDR < 0.05 cutoff (horizontal line).

A

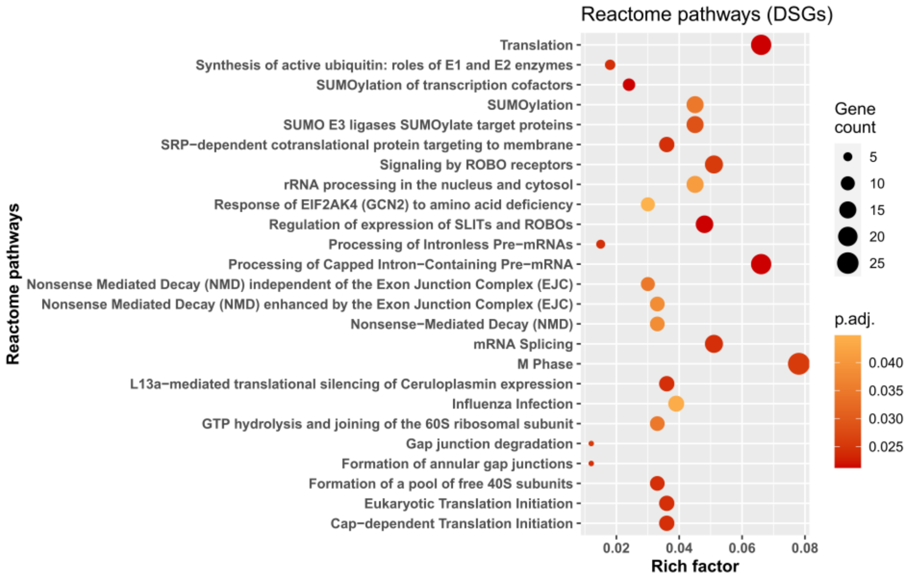


Supplemental Figure S3. Potential pathways regulated by DSGs detected upon *RBMXL3* overexpression and RNA-seq. (A) Reactome analysis for the DSGs based on the RNA-seq data. For the complete list of significant Reactome terms see Supplemental Table S6 (adjusted p-value < 0.05).

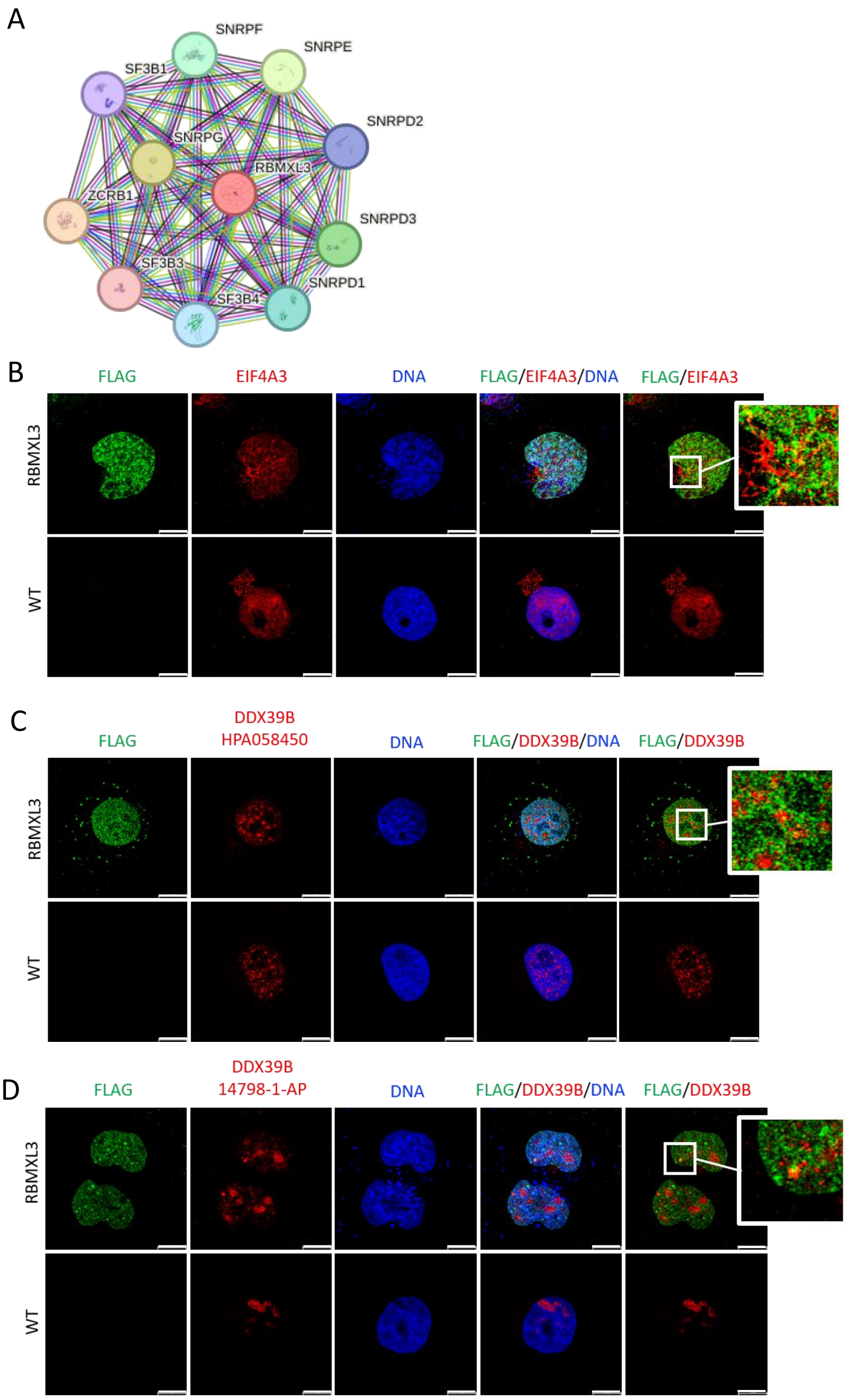


Supplemental Figure S4. STRING network for RBMXL3 and immunofluorescence co-staining of RBMXL3 and its potential protein interactors (EIF4A3 and DDX39B) identified by Co-IP-MS. (A) The STRING database network for RBMXL3 protein-protein interactions. (B) Confocal microscopy of TCam-2 cells expressing RBMXL3 and WT using anti-FLAG and anti-EIF4A3 antibodies. The zoom window illustrates signal colocalization of RBMXL3 and EIF4A3. Scale bar 11.6 µm. (C) Confocal microscopy of TCam-2 cells expressing RBMXL3 and WT using anti-FLAG and anti-DDX39B (HPA058450) antibodies. The zoom window illustrates signal colocalization of RBMXL3 and DDX39B. Scale bar 11.6 µm. (D) Confocal microscopy of TCam-2 cells expressing RBMXL3 and WT using anti-FLAG and anti-DDX39B (14798-1-AP) antibodies. The zoom window illustrates signal colocalization of RBMXL3 and DDX39B. Scale bar 11.6 µm.

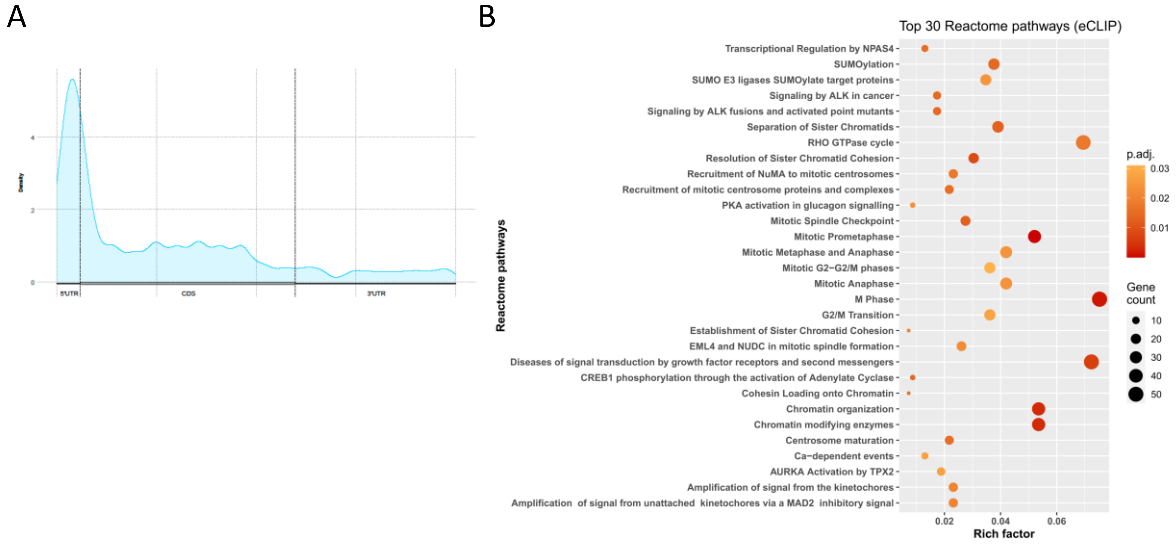


Supplemental Figure S5. Visualization of RBMXL3 binding and potential pathways regulated by RBMXL3. (A) Visualization of RBMXL3 RNA-binding sites with eCLIP peaks coverage alongside the standard mRNA model according to the MetaTX R package. (B) Reactome analysis for the transcripts bound by RBMXL3 based on eCLIP data. For the complete list of significant Reactome pathways see Supplemental Table S14 (adjusted p-value < 0.05).

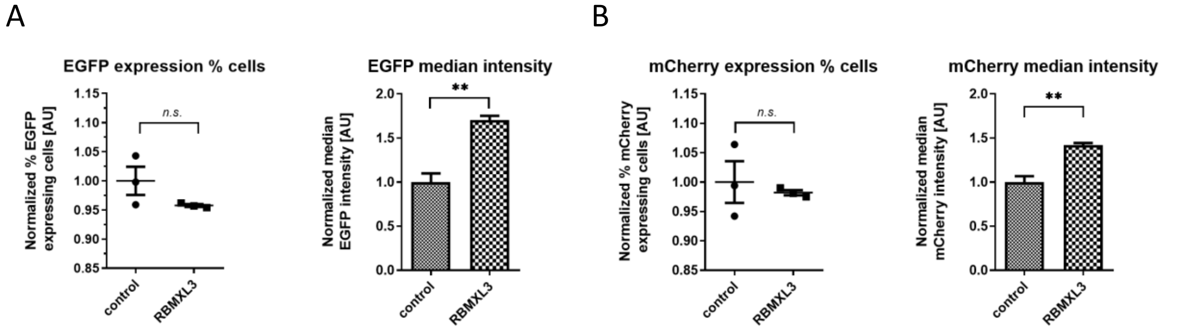


Supplemental Figure S6. LINE-1 retrotransposition assay control experiments show that RBMXL3 does not generally reduce EGFP and mCherry protein production which support the role of RBMXL3 in restricting the L1 retrotransposition. (A) Effect of overexpression of RBMXL3 and MBP (as control) on EGFP expression and EGFP median intensity in HEK293T cells. The results of independent experiments were normalized relative to the control (MBP). Statistical significance was calculated using Student's unpaired t-test (n.s. not significant; **$p < 0.01$; in comparison to MBP; each point = biological replicate). (B) Effect of overexpression of RBMXL3 and MBP (as control) on mCherry expression and mCherry median intensity in HEK293T cells. The results of independent experiments were normalized relative to the control (MBP). Statistical significance was calculated using Student's unpaired t-test (n.s. not significant, **$p < 0.01$; in comparison to MBP; each point = biological replicate). See Materials and Methods section for more details.
